# Virus-grazer interplay enhances virus production, particle aggregation, and trophic efficiency during infection of *Prochlorococcus*

**DOI:** 10.64898/2026.06.10.731369

**Authors:** Debbie Lindell, Michael C. G. Carlson, Julia Weissenbach, Shay Kirzner, Sigitas Šulčius, Yotam Hulata, Gazalah Sabehi, Svetlana Goldin, Joshua S. Sacks, Karin M. Björkman, Mathilde Dugenne, Sophia Zborowsky, Morgan D. Linney, Stephen J. Beckett, Ran Tahan, David Demory, Angelicque E. White, Joshua S. Weitz, David M. Karl, E. Virginia Armbrust, Anitra E. Ingalls, Francois Ribalet, David A. Caron

## Abstract

Viruses and grazers are fundamental agents of mortality in the oceans, impacting phytoplankton populations and organic matter cycling. Although viruses and grazers co-occur in nature, they are typically studied in isolation in laboratory experiments, limiting our understanding of their combined ecosystem impacts. Here, using a simplified ecosystem approach, we investigated individual and combined effects of the T7-like cyanopodovirus, P-SSP7, and the protistan grazer, *Paraphysomonas bandaiensis,* on the abundant marine cyanobacterium, *Prochlorococcus* MED4, and co-occurring non-photosynthetic heterotrophic bacteria (bacteria from here on). We observed that, individually, viruses and grazers caused substantial *Prochlorococcus* mortality. Viral lysis also triggered increases in damaged *Prochlorococcus* cells, dissolved organic matter release, and bacterial growth, while grazing reduced bacterial abundances. When grazers and viruses were combined, *Prochlorococcus* mortality was lower than expected from the sum of their individual effects. Contrary to expectations, this reduced *Prochlorococcus* mortality did not result in fewer viruses or grazers. Instead, virus-grazer-*Prochlorococcus* interplay resulted in greater virus production, maintenance of grazer growth, and a dramatic increase in particle aggregation. Our results reveal trophic cooperation and efficiency in which competition between viruses and grazers was likely mitigated, with virus progeny production enhanced by grazers, and grazer growth sustained through a shift to alternative food sources (bacteria, damaged cells, aggregates) secondarily derived from *Prochlorococcus* following viral lysis. The synergistic enhancement of particle aggregation via grazer-virus-phytoplankton interplay observed with the small buoyant *Prochlorococcus* phytoplankter underscores the importance of food web interactions for the flow of phytoplankton-fixed carbon within, and export from, the photic zone.

**Significance:** Viruses and grazers both use phytoplankton as a resource for reproduction. In a simplified experimental system with *Prochlorococcus*, an important primary producer in the oceans, we found that the interplay between viruses and grazers led to reduced mortality of *Prochlorococcus*. Despite this reduced mortality, virus-grazer interactions resulted in elevated virus production and a dramatic increase in organic matter aggregation. Furthermore, grazer abundance was not affected by this interplay, likely due to the transfer of organic matter from *Prochlorococcus* to bacteria and aggregates, which the grazers could consume as alternative food sources. These findings provide insights into the complexity of ecosystem interactions and how they impact the fate of organic matter fixed by phytoplankton in the oceans.

## Introduction

The oceans are teeming with microorganisms, forming a complex network of both mutualistic and antagonistic interactions. Phytoplankton are at the base of the food web and include a range of eukaryotic taxa and cyanobacteria, of which *Prochlorococcus* is the smallest and the most abundant globally (1). Ever present together with these primary producers are a diverse community of heterotrophic bacteria and archaea that are sustained largely on organic matter released from these autotrophs and other heterotrophic organisms (2–5). In turn, heterotrophic bacteria provide important goods and/or fulfill necessary functions that aid these autotrophs (6) as well as each other (7).

Phytoplankton also coexist in the water column with diverse agents of mortality whose antagonistic activities balance phytoplankton growth. These mortality agents include grazers that consume them and viruses that infect and lyse them. Grazers can generally feed on a wide range of organisms, with some degree of specificity acting at the level of prey size and/or taste (8, 9), although strain specificity has also been reported (10). Viruses are considerably more specific than grazers, infecting organisms within a particular taxonomic group, and even a specific genotype (11, 12). Grazers have also been shown to feed directly on viruses (13, 14) and can themselves be infected by viruses (15).

The process by which phytoplankton die affects the fate of photosynthetically fixed carbon. Lytic viruses kill their unicellular hosts releasing dissolved and particulate organic matter into the water column that can then be taken up and used by bacteria and archaea, stimulating their growth and affecting community composition (16–22). As such, viral lysis is important for recycling fixed carbon and nutrients in the photic zone. Viruses have also been implicated in stimulating export of organic matter as a result of particle stickiness and aggregate formation of lysed cells (23–26). Protistan grazing of phytoplankton results in the transfer of photosynthetically fixed carbon to higher trophic levels that subsequently lead to fecal pellet production by larger zooplankton and export from the photic zone through sinking and during diel vertical migration (27, 28). Sloppy feeding, egestion, and excretion by grazers can also release remineralized elements and organic matter that are utilized by other organisms within the photic zone (28–30). Thus, the mode of phytoplankton mortality is considered important in determining the partitioning of autotrophically fixed carbon, with viral lysis generally thought to contribute more substantially to recycling while grazing is thought to have a greater effect on carbon export, although both mortality agents contribute to each process (24, 28, 31).

Despite our growing understanding of the consequences of different mortality processes, little is known about the interplay between mortality agents (32, 33) and their combined ecosystem impacts since each mortality agent is typically studied in isolation. Here, using a simplified laboratory ecosystem approach we found a clear effect of the interplay between grazers and viruses. Despite reduced *Prochlorococcus* mortality, which is suggestive of competition between the grazer and the virus for this common resource, we observed elevated early virus progeny production and enhanced aggregate formation when grazers and viruses were present together, while grazer growth was not impacted. These findings highlight the complexity of interactions between food web components and establish the importance of such interactions for the flow of organic matter in the oceans.

## Results and Discussion

### Experimental set-up

In this study we investigated the individual and combined effects of virus infection and protistan grazing on cultures of *Prochlorococcus* MED4 and co-occurring non-photosynthetic bacteria (termed bacteria or heterotrophic bacteria from here on). As agents of mortality, we employed a single virus, the T7-like cyanopodovirus P-SSP7, that specifically infects *Prochlorococcus* MED4 (11), and a single grazer, the heterotrophic nanoflagellate *Paraphysomonas bandaiensis,* that is a voracious omnivore (34). The experimental design consisted of six main treatments (Fig. 1). *Prochlorococcus,* with an average starting abundance of 1.67±0.14×10^7^ cells·mL^-1^ (n=19), together with co-occurring heterotrophic bacteria, were grown with two initial concentrations of either grazers or viruses (Fig. 1a, Table S1) and compared to a control with just *Prochlorococcus* and the bacteria. The combined effects of both grazer and virus on *Prochlorococcus*, on the co-occurring heterotrophic bacteria, and on each other, were investigated using a treatment containing both low grazer and high virus concentrations (Fig. 1a, Table S1). The experiments were run in biological triplicates under a 14:10 light:dark cycle for 48 hours. The starting abundances of the organisms used, and the length of the experiment, allowed for two division cycles of *Prochlorococcus* in the control and for *Prochlorococcus* to decline and become a limiting resource for the mortality agents within a 12-48 h window as a result of grazing and viral lysis in the experimental treatments.

**Figure 1.**
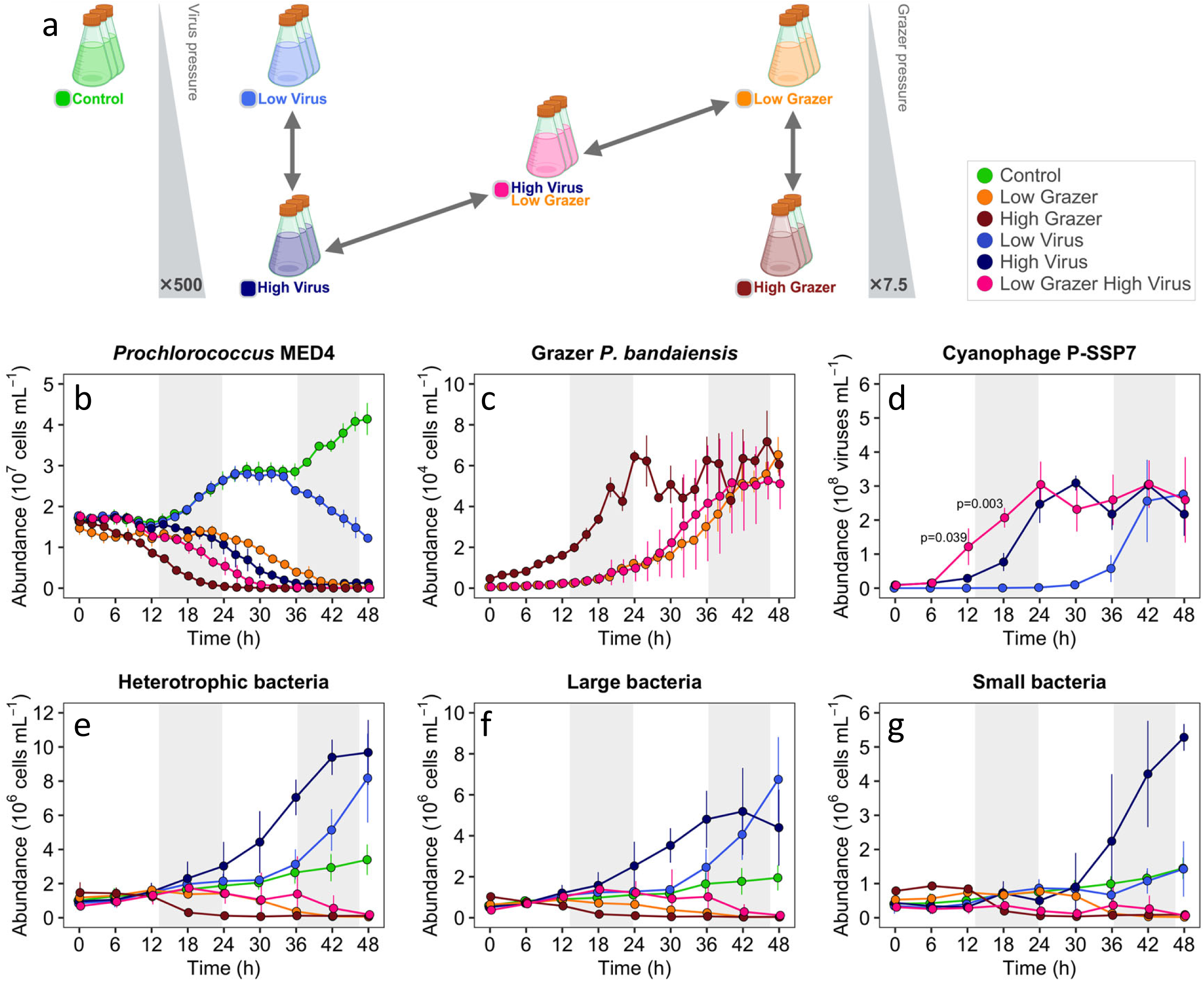
Experimental set-up and abundances of *Prochlorococcus* and mortality agents. Schematic representation of the experimental set-up (a). Treatments began with *Prochlorococcus* at an average of 1.67±0.14×10^7^ cells mL^-1^ without (control) or with low or high abundances of the cyanophage or grazer or both. Arrows indicate comparisons across treatments. All treatments were compared to the control of *Prochlorococcus* without grazers or viruses. Abundances of *Prochlorococcus* MED4 (b), the nanoflagellate grazer *Paraphysomonas bandaiensis* (c), the cyanophage P-SSP7 (d) as well as total (e), large (f) and small (g) heterotrophic bacteria. Average and standard deviation of 3 biological replicates. See Table S1 for starting abundances in the treatments. The number of viruses was significantly higher in the low-grazer-high-virus treatment than the high-virus treatment at 12 h (p=0.039) and 18 h (p-0.003), RM-ANOVA. See Table S2A,B,C,D,E,F for statistical analyses.

### Effects of virus and grazer on Prochlorococcus

*Prochlorococcus* grew exponentially in the absence of viruses and grazers, as determined from flow cytometric cell counts. The specific growth rate was 0.54±0.02 d^-1^ (n=4), doubling every 1.3±0.05 days. Growth was synchronized over the diel cycle with increased cell numbers at night (Fig. 1b) when cell division occurs and increased cell biomass during the day, as determined from particulate ATP concentrations (Fig. S1), when cells produce chemical energy and fix carbon via photosynthesis. These growth dynamics are similar to those expected for *Prochlorococcus* in nature (35–37).

The grazer exerted a large and rapid effect on *Prochlorococcus* populations. Exposure to initial low grazer concentrations (6.23±1.04×10^2^ grazers·mL^-1^, n=3) prevented an increase in *Prochlorococcus* abundance during the first night and resulted in a rapid decline during the second day (Fig. 1b). Exposure to 7.5-fold higher initial grazer concentrations (4.68±3.09×10^3^ grazers·mL^-1^, n=3) resulted in an immediate and rapid decline in *Prochlorococcus* abundances. The grazer population increased steadily over time (Fig. 1c) with a specific growth rate of 2.70±0.11 d^-1^, doubling every 6.2±0.25 h (n=6). The growth rate was similar in both the low and high grazer treatments during the exponential phase of growth. Grazer abundances reached a plateau of 5.4±1.01×10^4^ grazers·mL^-1^ (n=9) (Fig. 1c). This plateau was 10-fold less than the maximum grazer abundances we have observed when providing grazers with higher abundances of *Prochlorococcus* MED4 (see Materials and Methods), indicating that the grazer was resource limited at the plateau in these experiments.

*Prochlorococcus* numbers were negatively impacted by the virus. When exposed to a relatively low starting concentration of infectious viruses (2.4×10^4^ plaque-forming units per milliliter (PFU·mL^-1^), corresponding to a multiplicity of infection (MOI) of 0.0013), *Prochlorococcus* continued to grow over the first day of the experiment, both in biomass and numbers, with a clear decline relative to the control from 36 h onwards (Fig. 1b, Fig. S1). A 500-fold higher starting concentration of viruses in the high-virus treatment (1.2×10^7^ PFU·mL^-1^, MOI=0.7) resulted in a much more rapid decline in *Prochlorococcus,* apparent from 16 h onwards, in both cell numbers and particulate ATP concentrations (Fig. 1b, Fig. S1). Increases in virus abundances were apparent beginning at 30 h and 12 h in the low and high virus treatments, respectively (Fig. 1d), which was approximately 4-6 h prior to the observed declines in *Prochlorococcus* (Fig. 1b). *Prochlorococcus* lysis and virus progeny production occurred throughout the diel cycle as seen previously for this virus (38). The length of the lytic cycle was 6-9 h, as seen from the rise in virus release from cells during single-step growth experiments (Fig. S2). Virus abundances reached a plateau of 2.58±0.45×10^8^ viruses·mL^-1^ (n=9) (Fig. 1d), 3-10-fold less than those obtained on higher *Prochlorococcus* abundances (see Materials and Methods and Fig. S2), indicating that, similar to the grazer, the virus was resource limited once it reached a plateau in these experiments.

The combination of both low grazer and high virus resulted in a decline in *Prochlorococcus* abundance that was more rapid than observed for either of the mortality agents alone at the same initial abundances (Fig. 1b). However, the rate of *Prochlorococcus* decline in the combined treatment was less than the additive effect expected based on predictions from the individual treatments using a mortality model of functional responses to *Prochlorococcus* abundances (Fig. 2, Fig. S3). This less-than-additive effect could be the result of direct competition between the virus and the grazer for *Prochlorococcus* as a common resource, where consumption of *Prochlorococcus* by the grazer decreases *Prochlorococcus* availability to the virus, and vice versa, as has been proposed previously for host/prey in other settings (20, 32, 33). Alternatively, this could be due to multiple indirect mechanisms (Supplementary text), including direct grazer feeding on viruses which would reduce the number of viruses available for infection, a shift in grazer pressure away from healthy *Prochlorococcus* cells and towards other food sources in the system such as bacteria and dead *Prochlorococcus* cells, leaving more live cells available for use by both the grazer and the virus.

**Figure 2.**
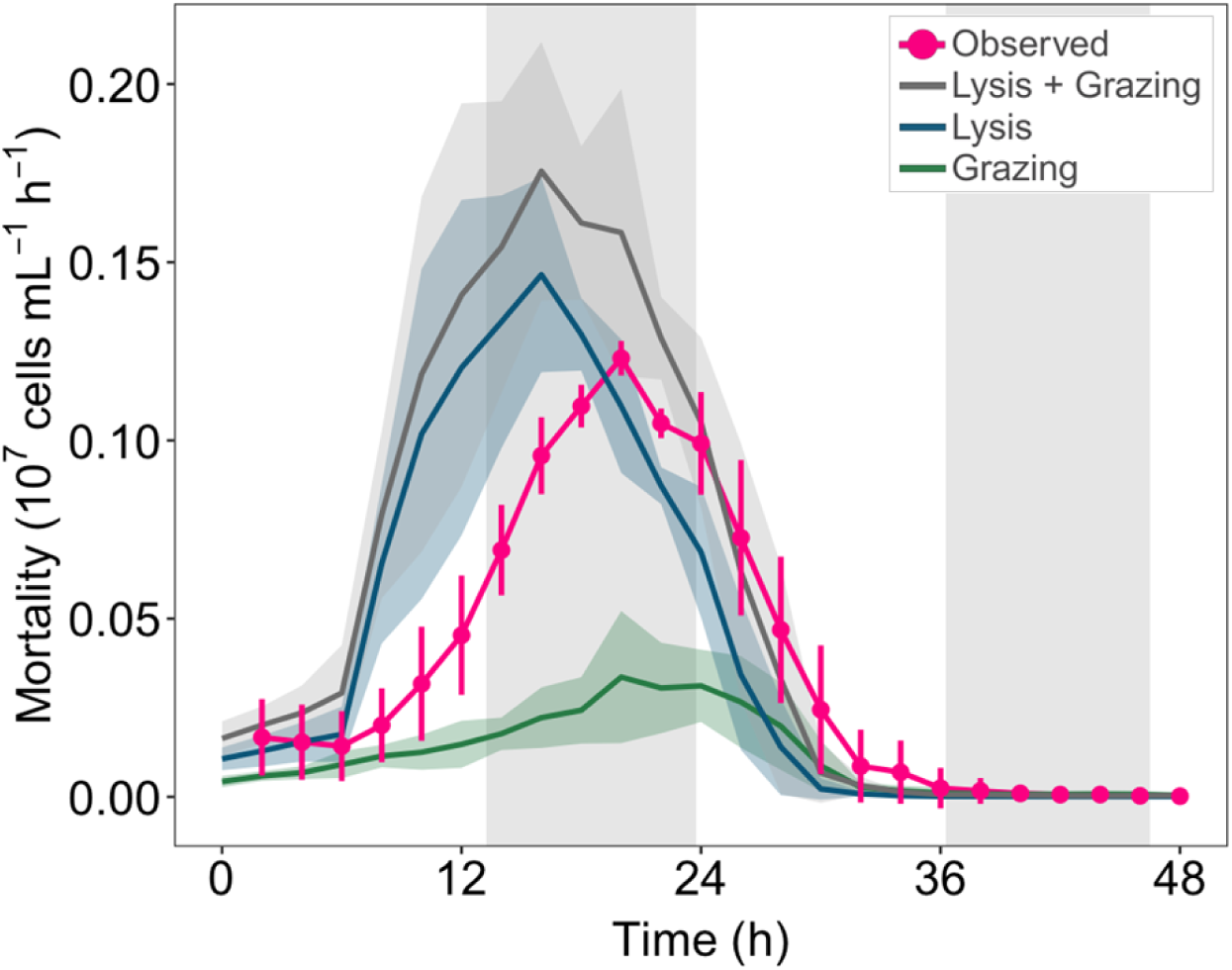
Predicted *Prochlorococcus* mortality in the presence of both virus and grazer. Predicted mortality of *Prochlorococcus* due to low grazer (green line), high virus (blue line), and both low grazer and high virus (grey line) relative to observed mortality in the low-grazer-high-virus treatment (pink line and circles). Observed mortality was calculated based on the difference in net growth rates between the control (assumed to have virtually no loss) and the treatments with low grazer and high virus. The predicted mortality for the combined low-grazer-high-virus treatment was calculated as the sum of the predicted loss rates induced by both grazer and virus, assuming independent and additive effects, and are based on a model that predicts grazing rates and viral lysis based on the functional responses of grazers and viruses to *Prochlorococcus* abundance. Vertical bars and the shaded area shows the standard deviation of 3 biological replicates. Vertical shading indicates nighttime. See Fig. S3 for mortality predictions relative to observed mortality of the individual treatments.

### Effects of virus and grazer on the bacterial community

Bacteria in the oceans grow on organic matter derived from multiple sources, including phytoplankton exudates during normal cellular growth (5), cells infected with and lysed by viruses (16–19, 31, 39, 22), and organic matter released from grazers and other bacteria (28–30). We measured select organic compounds that are typically found inside cells, such as glutamic acid (40, 41), ATP, and *Prochlorococcus* DNA in the dissolved phase (<0.2 µm) of the extracellular medium to provide an indication of released organic matter. In the *Prochlorococcus* control treatment (i.e. no mortality agent), these compounds were found in the extracellular medium (Fig. S4a). We observed considerably higher levels of dissolved organic matter for the treatments in which viruses were present (Fig. S4b,c,d, Table S2) (example for dissolved ATP at 24 h for high-virus: RM ANOVA p=0.0001; Tukey HSD p=1.1E-04), but not in the grazer-only treatment (Fig. S4e). These increases were transient in the high-virus and low-grazer-high-virus treatments (Fig. S4c,d), potentially due to use of these substrates by heterotrophic bacteria.

Heterotrophic bacteria were present at low levels in all treatments at the beginning of the experiments. Their abundances were, on average, 0.11±0.04×10^7^ cells mL^-1^ (n=19), which is 6.6% of initial *Prochlorococcus* numbers (Table S1). Bacterial abundances increased 3.5±1.3-fold (n=4) in the *Prochlorococcus* control cultures over the 48-h experiment (Fig. 1e), presumably growing on *Prochlorococcus* exudates. Bacterial abundances differed with treatment over the time course of the experiment (RM-ANOVA p=4.1E-29), with considerably more bacterial growth in the virus-only treatments (Fig. 1e), reaching 11.2±5.2-fold (n=6) higher numbers by the end of the experiment in both the low- and high-virus treatments (Tukey HSD, p<0.0001, n=6). In contrast, the presence of grazers, either alone or together with viruses, resulted in a significant decline in bacterial abundances (Fig. 1e). In the presence of both viruses and grazers, the decline was 4.5±1.0-fold (n=6) relative to the control (p<0.0001). In the grazer-only treatments this decline was considerably greater, 18.2±5.2-fold (n=6) relative to the control (p<0.0001) and occurred earlier (Fig. 1e). Thus, despite expected release of organic matter due to grazing that would stimulate bacterial growth (29, 30), a clear decline in bacteria was observed, likely due to strong grazing pressure on any bacteria that grew. The less dramatic decline in bacterial abundances in the combined virus and grazer treatment, despite the same number of grazers, was likely due to more substantial organic matter release as a result of virus-mediated cell lysis that stimulated more bacterial growth. These findings indicate that viral infection and lysis of *Prochlorococcus* enhanced bacterial growth, but that grazing dramatically diminished this stimulatory effect. Thus, different to competition between the grazer and virus for *Prochlorococcus* itself, virus-induced organic matter released upon lysis of *Prochlorococcus* enhanced bacterial growth, providing resources in another form to the grazer.

We discerned two different size classes of heterotrophic bacteria by flow cytometry in the virus treatments (Supplementary text). Proliferation of large bacteria (Fig. 1f) coincided with increases in virus numbers (Fig. 1d) and therefore with cell lysis, whereas growth of small bacteria (Fig. 1g) occurred once the plateaus in virus production (Fig. 1d) and large bacteria (Fig. 1f) were observed (Supplementary text). Large bacteria likely became resource limited whereas it appears that growth of the small bacteria was delayed either due to lack of suitable resources or the release of compounds from the large bacteria that inhibited their growth (Supplementary text). These results demonstrate that the two size classes of bacteria responded differently to *Prochlorococcus* viral lysis, resulting in succession between the large and small bacterial classes.

### The effects of virus and grazer on each other

Direct competition, and a number of potential indirect modes of interaction, between the virus and grazer, would be expected to negatively impact grazer reproduction and/or virus progeny production, and thus their abundances (32, 33). To assess whether such a negative effect exists, we compared virus and grazer abundances in the combined treatment (low-grazer-high-virus) to the treatments with either low grazers or high viruses. To our surprise, rather than a negative effect of the grazer on the virus, we found that virus abundances were higher in the presence of the grazer by 4.1±0.7-fold (n=3) and 2.9±0.9-fold (n=2) at 12 h and 18 h respectively (Fig. 1d) (RM-ANOVA p=0.008; Tukey HSD p=0.039 for 12 h and p=0.003 for 18 h). Once a plateau had been reached virus yields were similar in both the virus-only and grazer-virus treatments. Thus, virus abundances were not negatively impacted by the grazer and were, in fact, initially enhanced by their presence.

The temporal nature of the enhanced virus production led us to hypothesize that viruses were released earlier in the presence of the grazer. Indeed, we found that cellular compounds, such as glutamic acid and *Prochlorococcus* DNA were released to the extracellular medium earlier in the combined treatment (12-18 h) than in the virus-only treatment (18-24 h) (Fig. S4, Table S2). Furthermore, at 12 h we observed differential partitioning of virus DNA, with less virus DNA inside *Prochlorococcus* cells and more in the extracellular medium in the combined grazer-virus treatment relative to the virus-only treatment (Fig. 3, Fig. S5). Together these findings provide support for earlier cell lysis and virus release in the presence of the grazer. Such earlier release would allow a new cycle of infection to begin sooner and for overall greater production of viral DNA (Fig. 3, Fig. S5) and viruses (Fig. 1d) at 18 h in the presence of the grazer. Potential mechanisms leading to this early release (Supplementary text) include grazer-mediated changes in *Prochlorococcus* surface properties or physiology that could result in more rapid virus attachment to the cell or a shorter infection cycle, respectively. Alternatively, grazer ingestion of infected cells could result in early release of mature intracellular viruses since there is normally a lag between intracellular formation of viruses and their release from the cell.

**Figure 3.**
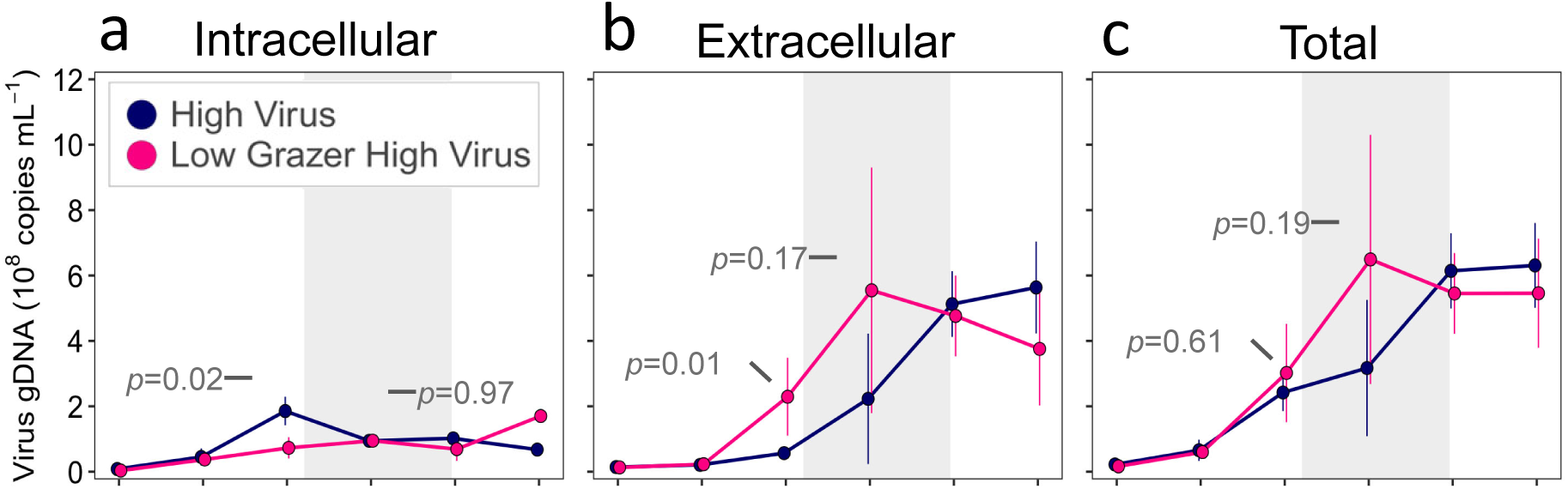
Partitioning of virus DNA in intracellular or extracellular fractions. (a) Intracellular virus DNA, (b) extracellular virus DNA, and (c) total virus DNA (intracellular + extracellular) during infection of *Prochlorococcus* MED4 in the absence (blue) and presence (pink) of the grazer. Average of n=3. See Fig. S5 for the individual replicates. Extracellular DNA includes encapsidated DNA packaged inside virus particles as well as unpackaged free DNA released from cells during lysis. Repeated measure analysis on log transformed data indicate a significant difference between the treatments. Post-hoc Tukey p-values for 12 and 18 h are shown. Note that while p-values were greater than 0.05 at 18 h for extracellular and total DNA, this is likely due to the variability in values between replicates since greater values were found in the combined treatment for all three replicates (Fig. S5). See Table S32I,J,K for statistical analyses.

In field settings, grazers have been reported to increase viral infection of bacteria and virus-like particle production (32, 42–44). This has generally been assumed to result from the indirect stimulation of the growth of the remaining bacteria that then serve as hosts for viruses. However, in our experiment the P-SSP7 cyanophage can only infect *Prochlorococcus* MED4 (11, 45), whose numbers declined rapidly in the presence of both grazer and virus. Thus, our findings indicate a direct effect of the grazer on viral production in *Prochlorococcus*, rather than through indirect stimulation of hosts for other viruses.

Next, we addressed whether the grazer was negatively impacted by the presence of the virus. Grazer abundances were similar in both the low-grazer-high-virus and the low-grazer treatments throughout the experiment (RM ANOVA; Tukey HSD p>0.1 across all time points; Table S2), with similar final yields (Fig. 1d) and specific growth rates (2.68±0.62 d^-1^ with the virus (n=3) versus 2.7±0.11 d^-1^ without the virus (n=6)). As such, no negative or stimulatory impact of the presence of viruses on grazers was observed, similar to previous findings in lake experiments (32).

The above findings raised the question of whether grazer populations were being maintained by acquiring carbon and energy from sources other than healthy *Prochlorococcus* cells. In contrast to certain grazer-virus pairs (13, 14, 46), we found that this grazer did not directly feed on this virus (Fig. S6, Supplementary text). Another food source for the grazer, as described above, was likely to be the co-occurring heterotrophic bacteria whose growth was stimulated by organic matter released from lysed *Prochlorococcus* cells (Fig. 1e). Furthermore, grazers could obtain additional resources by feeding on infected and damaged cells resulting from viral lysis as well as on aggregates and cell debris that were generated in the combined grazer-virus treatment (see below). Thus, maintenance of grazer growth and yields in the presence of the virus was likely facilitated through grazer recapture of organic matter derived from *Prochlorococcus* as the result of viral lysis.

### Effects of virus and grazer on Prochlorococcus cell damage and particle aggregation

Changes in particle size structure and chlorophyll fluorescence developed over the course of the experiment, providing insight into the transformation and fate of *Prochlorococcus* as a function of the mortality processes it was subjected to. Three fluorescent flow cytometric populations were evident in addition to the healthy *Prochlorococcus* population (Fig. 4a). One such population had similar chlorophyll fluorescence to the healthy *Prochlorococcus* population, but less forward scatter (Fig. 4a, leftmost population in blue in all panels; low-scatter population), and thus potentially consisted of damaged *Prochlorococcus* cells. The second fluorescent population displayed greater forward scatter than healthy *Prochlorococcus* populations and a range of fluorescence levels (Fig. 4a, to the right of *Prochlorococcus* in red in all panels; intermediate-scatter population) and thus could consist of aggregates. The third fluorescent population had the highest forward scatter (Fig. 4a, far right in purple in panels iii and iv; high-scatter population) and was observed only in the grazer treatments and thus could be grazers that have ingested *Prochlorococcus* cells. These cytometric populations likely maintained their fluorescence since chlorophyll can be stable for days (47, 48). We quantified these populations and found that they displayed different relative abundances depending on the experimental treatments (Fig. 4, Fig. S7). We also sorted and imaged these fluorescent populations by scanning electron microscopy (SEM) to visually characterize them (Fig. 4, Fig. S8,S9,S10).

**Figure 4.**
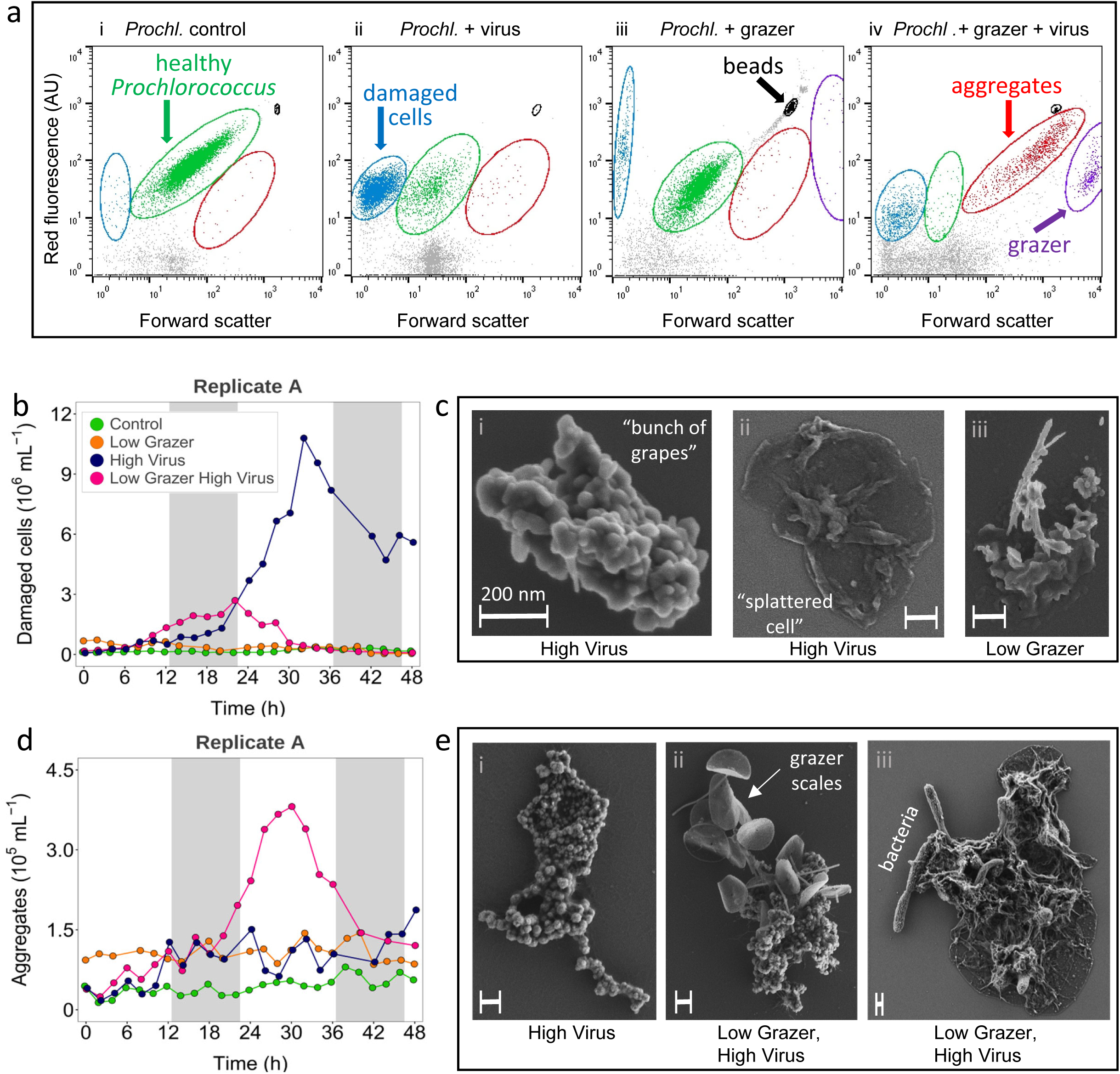
Appearance of damaged *Prochlorococcus* cells and fluorescent aggregates. Representative cytograms (a) showing the healthy *Prochlorococcus* population (green), damaged cells (low-scatter population in blue), fluorescent aggregates (intermediate-scatter population in red) and grazers with *Prochlorococcus* accumulation (high-scatter population in purple) at T=36 h in the *Prochlorococcus* control (i), *Prochlorococcus* + high virus (ii), *Prochlorococcus* + low grazer (iii), and *Prochlorococcus* + low grazer + high virus (iv) treatments. Beads are shown in black. AU – arbitrary units. Representative graphs of abundance (b,d) and SEM images (c,e) of damaged *Prochlorococcus* cells (b,c) and fluorescent aggregates (d,e). SEM images are from T=32 h. The treatment source is shown below each image (see Table S4 for more details). The scale bar, shown in white, is 200 nm in length in all images. See Fig. S7 for quantification of damaged cells and aggregates for all 3 replicates and Figs. S9 and S10 for additional SEM images of damaged cells and fluorescent aggregates, respectively. See Table S3 for timing of treatment-specific changes in damaged cells and fluorescent aggregates, and Table S2G,H for statistical analyses.

The low-scatter population (Fig. 4a, blue) was found in all treatments. It was particularly prevalent in the high-virus treatment with an increase that ranged from 18 to 35-fold beginning at 12-18 h and that peaked at 28-34 h (Fig. 4b, Fig. S7a). This paralleled the decline in the number of healthy *Prochlorococcus* cells and the increase in virus abundances (Fig. 1b,d). The low-scatter population was also enriched in the combined low-grazer-high-virus treatment, although significantly less so than in the high-virus treatment (1.6-4.2-fold less, Kruskal-Wallis p=4.0E-07, n=15) (Fig. 4a,b, Fig. S7). The abundance of this population declined with time in both virus-containing treatments (Fig. 4b, Fig. S7a,b), contrasting with the control where the low-scatter population increased 2-5-fold by the end of the experiment (Fig. S7b) (Wilcoxon test, p=5.8E-07, n=3). These phenomena were consistently observed for all three biological replicates, but the timing of their appearance and magnitude of change varied (Fig. S7, Table S3).

Relative to the healthy *Prochlorococcus* population which consisted of coccoid cells approximately 500-700 nm in diameter and dividing cells (Fig. S8a), SEM images of the low-scatter population showed a variety of misshapen cells and other cellular debris (Fig. 4c, Fig. S9). Prevalent forms visually resembled “bunches of grapes” (image (i) in Fig. 4c and Fig S9a,b,c) and “splattered cells” (image (ii) in Fig. 4c and Fig. S9a,b,c) as well as more diffuse cellular debris in various forms (image (iv) in Fig. S9a,b,c). The latter diffuse type, but not the other types, was also observed in the low-grazer treatment (Fig. S9d). These images support our hypothesis that the high abundance of this low-scatter cytometric population in the treatments with viruses was likely due to damage caused by virus-mediated cell lysis.

Large amounts of submicron organic matter, such as those exemplified by the low-scatter population, are known to be suspended in the upper layers of the oceans (49, 50). Our findings, together with those of Shibata et al. (51), who report similar production of submicron particles for a *Vibrio*-virus marine system, suggest that viral lysis is likely to be a major source of such suspended organic matter.

Abundance of the intermediate-scatter chlorophyll-fluorescing cytometric population (Fig. 4a, red) also depended on the treatment. This population was observed in all treatments with a mild increase of 1.1-1.8-fold in the control (Wilcoxon test, p=0.003, n=4) and a more substantial increase of 1.9-4.1-fold in the high-virus treatment over time (Fig. 4d, T-test, p=5.3E-10, n=3). This intermediate-scatter population was particularly enhanced in the low-grazer-high-virus treatment for a 10-16 h period in the middle of the experiment (7.3-10.7-fold relative to the beginning of the experiment), significantly more so than in the high-virus (Kruskal-Wallis H test; p=8.1E-09, n=15) or low-grazer treatments (Fig. 4d, Fig. S7c). The appearance of this population was consistent across replicates (Fig. S7c) and peaked 4-8 h after the peak in damaged cells in each replicate (Figs. S7a,c).

SEM imaging of the intermediate-scatter fluorescent population showed that it consisted of aggregates. The aggregates observed in our cytogram window ranged in size from approximately 2 to 10 µm and contained *Prochlorococcus* cells, other bacteria, grazer scales and cellular debris of various types including the “bunches of grapes” (images i and ii) and “splattered cells” (image iii) forms (Fig. 4e, Fig. S10). The coincidence of aggregate formation with the decline in damaged cells in each replicate of the low-grazer-high-virus treatment (Fig. 4, Fig. S7) together with the observation of damaged cell forms in aggregates, suggests that the cell debris produced as a result of viral lysis became an integral part of these aggregates. Interestingly, damaged cells in the forms of “bunches of grapes” and “splattered cells” were observed in low amounts in the control treatment aggregates (Fig. S10a), but were not observed in the damaged cells or aggregates from the low-grazer treatment (Fig. S9c, Fig. S10d), raising the possibility that they were fed upon by the grazer.

The prevalence of damaged cells and aggregates declined in the latter part of the experiment (Fig. 4, Fig. S7). This could be due to continued aggregation to larger sizes, loss of fluorescence over time, feeding by the grazer (when present), or a combination of these processes. Disaggregation and bacterial degradation of the aggregates to smaller sizes seem unlikely, since no new appearance of small particles occurred.

Grazers and viruses are known to individually result in aggregated organic matter from their hosts and prey in other systems (26, 52–54). Our findings indicate that viral infection of phytoplankton as small as *Prochlorococcus* can also result in organic matter aggregation. This is even though uninfected *Prochlorococcus* was neutrally buoyant and that infected *Prochlorococcus* cells were positively buoyant (Fig. S11, Supplementary text). This aggregation could be partly due to the release of sticky organic material from *Prochlorococcus* itself under the stress of infection or produced from co-occurring bacteria (55, 56) whose growth was stimulated as a result of cell lysis (Fig. 1e). We propose that the dramatic increase in particle aggregation observed in the combined grazer-virus treatment resulted from a synergistic effect of the release of organic material from *Prochlorococcus* after viral lysis and/or other bacteria that grew subsequent to this lysis, and from the protistan grazer via egestion during feeding.

The high-scatter population was evident only in treatments with the grazer (Fig. 4a) and consisted of grazers as seen from SEM images (Fig. S8b). The flagellum was often obvious (Fig. S8b images i-iv), as was the presence of signature scales for *P. bandaiensis* (Fig. S8b images iii-iv, vii-ix). A flimsy sheet-like structure was also often observed (Fig. S8b images vii-ix). We propose that actively feeding grazers had ingested sufficient numbers of *Prochlorococcus* whose chlorophyll had not yet degraded inside the food vacuole (57), hence their chlorophyll fluorescence. Sometimes aggregates were attached to the grazer cell, particularly in the combined grazer and virus treatment (Fig. S8b images v-ix). This could result from material egested from food vacuoles that became stuck to the grazer (58) or their attachment could be indicative of grazers feeding on the aggregates.

### Ecosystem consequences of the interplay between the grazer and the virus

Our simplified laboratory ecosystem experiments revealed multiple consequences of the interplay between grazers and viruses. The most striking of these was the dramatic increase in aggregation of particulate matter far surpassing that for either grazers or viruses alone. These aggregates were formed from a combination of cellular debris produced as a result of viral lysis, as well as from *Prochlorococcus* cells (whether healthy, lysed by the virus, or egested from the grazer), heterotrophic bacteria, and grazer structures. Virus-induced aggregation and export from the photic zone was initially thought to occur primarily for ballasted phytoplankton such as coccolithophores and diatoms (25, 26). However, findings of DNA of both cyanophages and picocyanobacteria in deep waters and sediment traps have led to the suggestion that viral lysis of smaller buoyant phytoplankton can also lead to cellular aggregation and sinking (23, 59). Our findings provide empirical support for this suggestion and reveal a synergistic enhancement of particle aggregation as a result of the interaction between viral lysis and protistan grazing on *Prochlorococcus*.

The fate of aggregates in the oceans has great importance for carbon cycling. Aggregates serve as an important substrate for bacterial colonization and remineralization of the organic matter contained within (60, 61). Small aggregates such as those observed in our study could be suspended or sinking (62). Even if suspended it seems probable that they could further agglomerate to form larger and larger particles that could sink out of the photic zone (27, 62, 63). These aggregates, and the bacteria within them, could also be fed upon by protists and other zooplankton (27, 28, 64–66) moving some organic matter up the food chain to be eventually released from larger zooplankton in fecal pellets. Thus, irrespective of the pathway, aggregation due to the interplay between viruses and grazers likely enhances the flux of organic matter out of the photic zone as part of the biological pump.

Another unexpected consequence of grazer-virus interplay is the absence of a negative effect on grazer growth or virus progeny production. This was despite resources becoming limiting in the system, as observed from plateaus in grazer and virus abundances (Fig. 1c,d), and lower than expected *Prochlorococcus* mortality in the combined grazer-virus treatment (Fig. 2). In particular, the positive direct effect of the grazer on early virus production was surprising. These findings suggest that in an open ecosystem, where the shared resource does not end as it did in our experimental system, viral infection would be higher than if such an interplay did not exist. Grazers were also not disadvantaged by their interaction with viruses. We propose that this is due to the grazer’s omnivorous nature, allowing it to shift to feeding on organic resources that were transformed from *Prochlorococcus* to other forms as a result of viral lysis and grazer-virus-*Prochlorococcus* interplay, such as the bacteria that grew subsequent to cell lysis, as well as the damaged cells and aggregates that developed. Such recouping of organic matter secondarily derived from *Prochlorococcus* could also serve to somewhat diminish grazing pressure on healthy *Prochlorococcus* populations. In this scenario there is very little waste, with the live *Prochlorococcus* cell providing the virus with the resources needed to produce new virions, the organic matter released from the lysed cell fueling bacterial growth, and the damaged cell remains being fed upon by the grazer either directly or in aggregates. Our results showing the reciprocal benefit of the grazer on early virus production and the virus on the supply of alternative food sources for the grazer, suggest that while grazers and viruses compete for common autotrophic resources, the evolution of traits among microbes within ocean ecosystems has led to the emergence of a type of trophic cooperation and trophic efficiency that mitigates this competition.

## Materials and Methods

### Strains and culture conditions

*Prochlorococcus* MED4, a unicellular cyanobacterium belonging to the HLI clade was the cyanobacterium used in this study. Xenic cultures were grown in a seawater based Pro99 medium (67) with bicarbonate (12 mM) but without an additional buffer. Cultures were incubated at 21 °C with very slow stirring under a 14:10 light:dark cycle with 50 µmol·photon·m^-2^·s^-1^ cool white light during the light period. Three independent cultures of *Prochlorococcus* were maintained in exponential growth for multiple generations prior to experimentation. The cyanophage, P-SSP7, a T7-like podovirus that infects *Prochlorococcus* MED4 (11), was propagated on exponentially growing host cells under the above growth conditions, reaching an infective titer of 3×10^9^ phages·mL^-1^. The grazer, *Paraphysomonas bandaiensis*, a heterotrophic, phagotrophic, nanoflagellate belonging to the chrysophytes (34), was initially grown in a seawater medium containing 10% yeast extract broth with the addition of rice grains to stimulate bacterial growth for the grazer to feed upon (34). The grazer was transferred to a diet of *Prochlorococcus* MED4 in Pro99 medium and maintained in exponential growth for multiple generations until the beginning of the experiments. Maximal yields of the grazer on a diet of *Prochlorococcus* reached 6.0±1.8×10^5^ grazers·mL^-1^.

### Experimental design

The six main experimental treatments consisted of a control culture of *Prochlorococcus* with no mortality agent, treatments with low or high concentrations of grazer or virus and a combined treatment with a high concentration of viruses and a low concentration of grazers (Fig. 1, Table S1). Additional treatments consisted of grazers and viruses, or just grazers, without *Prochlorococcus*, and a high-grazer-high-virus treatment.

Cultures of *Prochlorococcus* MED4 were initially grown to a concentration of 1.6-1.8×10^8^ cells·mL^-1^ in exponential phase. At the beginning of the experiment, cultures were diluted 10-fold into fresh medium and grown in 10 L polycarbonate carboys (Nalgene) under the conditions described above. Exponentially growing grazers were added at average abundances of 600-620 grazers·mL^-1^ and 4700 grazers·mL^-1^ for the low and high treatments respectively. Viruses were added at ratios of 0.0013 and 0.7 infective viruses to *Prochlorococcus* cells (MOI) for the low and high virus treatments, respectively, and allowed to adsorb to *Prochlorococcus* for 90 min (on average) prior to the above-mentioned 10-fold dilution of the cultures at the beginning of the experiment. This corresponded to a beginning concentration of 1.71x10^7^ *Prochlorococcus*·mL^-1^ with 2.4×0^4^ and 1.2×10^7^ infective viruses·mL^-1^ added to the low and high virus treatments respectively. Cultures of *Prochlorococcus* and the grazer were not axenic and contained heterotrophic bacteria ranging from 0.68-1.47x10^6^ cells·mL^-1^ at the beginning of the experiments (Table S1). This is between 4-9% of the *Prochlorococcus* abundances. Initial abundances of the mortality agents were determined empirically in preliminary experiments so that growth and decline of *Prochlorococcus* MED4 occurred within a 12-48 h window. All treatments were carried out in biological triplicates. Growth rates of *Prochlorococcus,* grazers and bacteria were determined for the exponential phase of growth.

### Sampling and analyses overview

The first samples (T=0) were collected immediately after dilution of *Prochlorococcus*, with the addition of the mortality agents when present, 1.5 h after the light cycle started, and at the time intervals described below over a 48-h period. Samples were collected every 2 h for the determination of *Prochlorococcus* abundances based on their red autofluorescence and forward scatter by flow cytometry. These same samples were used for enumeration of damaged cells and fluorescent aggregates. They were also used for flow cytometric sorting into four discrete fluorescent populations based on red autofluorescence and forward scatter for SEM imaging. Grazer abundances were determined every 2 h and heterotrophic bacteria every 6 h by flow cytometry after staining with a DNA stain. Samples were collected every 6 h for the determination of virus abundances by quantitative PCR (qPCR) after DNase treatment to remove unencapsidated DNA. Intracellular and extracellular virus and *Prochlorococcus* DNA were also quantified every 6 h by qPCR. Samples were collected every 12 h for quantification of organic matter in the extracellular medium; glutamic acid was measured using a cation-exchange solid phase extraction paired with liquid-chromatography-mass spectrometry, particulate and dissolved ATP was measured using the firefly bioluminescence reaction, and overall DNA concentrations were determined using a fluorescence assay. See below and Supplementary methods for details of all procedures.

### Quantification by flow cytometry

Flow cytometry samples (1 mL) were fixed in 0.1 % glutaraldehyde for 15 min in the dark at room temperature, flash frozen in liquid nitrogen and stored at -80°C prior to analyses. An Influx flow cytometer (BD Biosciences), equipped with a small-particle forward scatter detector and a 70 µm nozzle, was used for quantification and sorting. *Prochlorococcus* was detected based on its chlorophyll red autofluorescence (using a 692/40 nm bandpass filter) and characteristic forward scatter as a proxy for size when excited with both a 488 nm and a 457 nm laser. Data collection was triggered by the 488 nm scatter signal. The red autofluorescence runs were also used to quantify damaged *Prochlorococcus* cells and red fluorescing aggregates, and to detect grazers with sufficient chlorophyll inside their vacuoles after ingestion of *Prochlorococcus*. Flow cytometric populations were differentiated from *Prochlorococcus* and each other based on their forward scatter (Fig. 4a). These same cytometric populations were sorted for scanning electron microscopy imaging.

Heterotrophic bacteria and grazers were stained with a 10,000-fold dilution of the SYBR™ Green I dsDNA stain (Invitrogen). Total bacteria and grazers were detected based on the green fluorescence of the DNA stain (using a 530/40 nm bandpass filter) and their forward scatter after excitation with the 488 nm laser. To differentiate between *Prochlorococcus* and heterotrophic bacteria in the stained runs, data were analyzed with R software using the FCSplankton library version 1.1.0. *Prochlorococcus* was discriminated from heterotrophic bacteria based on its dual fluorescence in the red (autofluorescence) and the green (from the DNA stain). Total heterotrophic bacteria (considered to be all bacteria without chlorophyll fluorescence) were determined from the difference between total bacteria and *Prochlorococcus* abundances. “Small” and “large” heterotrophic bacteria were discriminated from each other based on their light scatter and green fluorescence, with the large bacteria having both more forward scatter and higher green fluorescence.

The photomultiplier tube for forward scatter of the 488 nm laser was set at 27 V for *Prochlorococcus* autofluorescence runs, at 36 V for total bacterial counts and 16 V for grazer counts. All runs included 1 µm FluoSphere carboxylate yellow-green fluorescent beads (Invitrogen) which served as an internal standard for fluorescence and forward scatter.

### Virus quantification

Samples were filtered through a 0.22 µm PVDF syringe filter (Millex-GV, Millipore) and the filtrate was treated with 5 U of DNase I for 60 min at 37°C to degrade all unencapsidated DNA followed by inactivation of the enzyme with 50 mM EDTA (pH8) (68). The filtrates were diluted 100-fold in 10 mM Tris buffer (pH8) to eliminate inhibitory effects of seawater on PCR. Duplicate SYBR Green qPCR assays (LightCycler® 480 SYBR Green I Master Mix from Roche Diagnostics) were used to quantify encapsidated virus DNA. Specific primers (0.5 µM) for the P-SSP7 cyanophage DNA polymerase gene were used ((Forward: AACACTTCCGCCCTTACCT and Reverse: CTGCAACGAAAGGGAATTGT) (69)). Thermocycling consisted of an initial denaturation step of 10 min at 95°C, and 40 cycles of denaturation at 95°C for 10 s, annealing at 57°C for 10 s, and elongation at 72°C for 10 s. Fluorescence plate read measurements were carried out at the end of each cycle. Melting curve analysis from 40 to 97°C with a fluorescence read each 1°C verified the specificity of the amplified products. Standard curves of a dilution series of known concentrations of the P-SSP7 cyanophage (based on virus-like particle quantification (70)) were used to quantify virus abundances.

### Scanning electron microscopy

Sorted cytometric populations in 25 µL volumes were placed on poly-L-lysine coated silicon chips. The samples were fixed in 2% glutaraldehyde and 2% paraformaldehyde in a 0.1 M cacodylate buffer (pH4). The samples were post-fixed using 1% osmium tetra-oxide, dehydrated in a dilution series of ethanol from 10 to 100%, critical point dried (Quorum K850), and coated with a ∼2 nm layer of graphite (Q150T ES Plus Coater). Samples then were viewed on a Zeiss Ultra Plus HR Scanning Electron Microscope using a Secondary Electron detector and an In-lens detector.

### Prochlorococcus mortality estimates

*Prochlorococcus* treatment-induced mortality (grazing, viral lysis, or both grazing and lysis) was estimated by comparing net population growth rates between control and experimental treatments, assuming per-capita cell division was unchanged across treatments. Net growth rates were calculated as the slope of the natural log of (smoothed) *Prochlorococcus* abundance versus time. Mortality attributable to a treatment was computed as the difference between control and treatment net growth rates. To relate prey abundance to loss processes, per-capita viral lysis and grazing rates were modeled as functions of *Prochlorococcus* abundance using a Holling type I functional response (linear) and Holling type II functional response (hyperbolic), respectively (71) (Fig. S12 and Supplemental text). Fitted models were used to predict loss rates under each condition, scaled by grazer or virus abundance. Expected mortality in the combined grazer-virus treatment was calculated as the sum of predicted grazing and viral losses (additive effects), following multitrophic *Prochlorococcus* mortality modeling approaches (72). See Supplemental methods for more details. The data and code used for modelling are available on Github (https://github.com/fribalet/PROMO_modelling).

### Statistical analyses

All variables were screened for whether they satisfied the assumptions of normality (Shapiro-Wilk W-test and Q-Q plots), sphericity (Mauchly’s Test) and homoscedasticity (Levene’s test) required for parametric tests. Data were log-transformed when the Shapiro-Wilk test was significant (p<0.05) and visual inspection of the Q-Q plots confirmed asymmetry. Greenhouse-Geisser corrections were applied whenever Mauchly’s test indicated a violation of sphericity (p<0.05). Differences in abundances, intra- and extracellular DNA and metabolite concentration measurements over the course of the experiments were assessed by using a repeated-measures analysis of variance (RM-ANOVA), with “time” as within-subjects and “treatment” as between-subjects (independent) factors. When results were significant, Tukey’s post hoc multiple pairwise comparisons were performed to determine where and when significant differences occurred between the different treatments. For cases of unequal sample sizes (e.g. missing values) and/or when conditions for equal variances (homoscedasticity and sphericity) were not met, linear mixed effects regression was used. We used a combination of Rodionov’s STARS (73), Bayesian (74), and PELT (75) change point analyses to detect either gradual or abrupt differences (if any) in timing of the treatment-specific effects on damaged *Prochlorococcus* cells and fluorescent aggregates, followed by ANOVA or Kruskal-Wallis rank sum test to determine whether those effects differ between the treatments. All results were considered significant at p≤0.05.

## Acknowledgements

We thank Tara Clemente, Sean Kearney, Dror Shitrit, and Ilia Maidanik for help with sampling, Lihi Shaulov from the Electron Microscopy unit at the Biomedical Core Facilities, Faculty of Medicine, Technion, for SEM sample preparation and Maria Koifman Khristosov from the Electron Microscopy Center, Department of Materials Science and Engineering, Technion, for SEM imaging, Everetta Rasyid and Laura Carlson for laboratory assistance with glutamic acid analyses. We thank SCOPE participants and Lindell lab members for discussions. This research was funded by the Simons Foundation (SCOPE grants 329108 and 721254, Life Sciences grant 529554 and SFI-LS-PROJECT-11154 to DL, SCOPE grant 721256 to AEW, SCOPE grant 721231 to JSW, SCOPE grant 329108 and Life Sciences grant SFI-LS-PROJECT-9452 to DMK, SCOPE grant 721244 and Life Sciences grant SFI-LS-PROJECT-12202 to EVA, SCOPE grant 329108 and Life Sciences grant 385428 to AEI, Life Sciences grant 574495 to FR, SCOPE grant P4802 to DAC). SJB and JSW are investigators at the University of Maryland-Institute for Health Computing, which is supported by funding from Montgomery County, Maryland and The University of Maryland Strategic Partnership: MPowering the State, a formal collaboration between the University of Maryland, College Park and the University of Maryland, Baltimore.

## Author contributions

DL, JSW, FR, DAC conceived the project, DL, MCGC, JW, GS designed the experiments with contributions from all authors, DL, MCGC, JW, SK, SS, YH, GS, SG, JSS, KMB, MD, SZ, MDL, SJB, RT, DD carried out the experiments, DL, MCGC, JW, SK, SS, YH, GS, SG, JSS, KMB, MD, SZ, MDL, RT, DD, DMK, FR analyzed the data, DL, AEW, JSW, DMK, EVA, AEI, FR supervised the research, DL wrote the first draft of the manuscript and all authors contributed intellectually to the research and to revisions of the manuscript.

## Supplementary Text: Supplementary Results and Discussion

### Direct or indirect competition for Prochlorococcus

The less-than-additive impact of viruses and grazers on *Prochlorococcus* mortality could be the result of direct competition for *Prochlorococcus* as a common resource. However, this effect could also be, at least partially, indirect and due to one or more possible processes: Grazer feeding on the virus or infected cells that would reduce the number of viruses available for infection (1); a physiological response of *Prochlorococcus* that affects infection and/or grazing and thus mortality; physical interference by grazer- or virus-released materials that prevents access of the other to *Prochlorococcus;* a shift in grazer pressure away from *Prochlorococcus* and towards other food sources in the system including heterotrophic bacteria, lysed cells and aggregates; double use of the same cells for both virus production and grazer nutrition that would leave more live *Prochlorococcus* cells for use by both the virus and the grazer.

### Virus-induced lysis stimulated growth of two different size classes of bacteria

We discerned two different size classes of heterotrophic bacteria in the virus treatments by flow cytometry. The large bacteria had greater forward scatter and more DNA fluorescence (after staining) than the small bacteria. In both the low- and high-virus treatments, growth of large bacteria was observed earlier than growth of small bacteria (Fig. 1f,g). Proliferation of the large bacteria coincided with the increase in virus numbers and therefore with cell lysis, after 12 h in the high-virus treatment and 30 h in the low-virus treatment (Fig. 1f). The net growth rate of the large bacterial consortium at the steepest incline was similar in both the high- and low-virus treatments, doubling every 8.6±0.9 h (n=6). A plateau in the abundance of the large bacteria was observed in the high-virus treatment from 36 h. At this time the small bacteria increased in abundance (Fig. 1g) and had a growth rate approximately two-fold faster than that of the large bacteria, doubling every 4.6±0.6 h, (n=3). The plateau in large bacteria and growth of the small bacteria occurred within the period of the plateau in virus production (Fig. 1d) and coincided with both the leveling off in the decline of *Prochlorococcus* (Fig. 1b) and the drawdown of dissolved organic matter (Fig. S4c). By contrast, in the low-virus treatment a plateau in large bacterial abundances was not reached and there was no substantial growth of small bacteria (Fig. 1f,g). Correspondingly, *Prochlorococcus* abundances continued to decline (Fig. 1b) and the concentration of dissolved glutamic acid and ATP continued to rise (Fig. S4b). These combined findings suggest that the large bacteria became resource limited upon the decline in the supply of organic matter released from *Prochlorococcus* due to viral lysis and that growth of the small bacteria was delayed until this time.

The delay in growth of the small bacteria is intriguing. This could be due to differences in substrate specificity of the members of the two bacterial size classes (2), utilizing different organic compounds released from *Prochlorococcus* at different stages. Indeed, Sacks et al. (3) found a change in the dominant metabolites in *Prochlorococcus* in the presence of viruses towards the end of the experiment which would affect the metabolites released upon cell lysis. Alternatively, it’s possible that the small bacteria utilize compounds exuded from the large bacteria at stationary phase or that some of the large bacteria produce compounds during growth that inhibit growth of the small bacteria. While the mechanism for the delay in growth of the small bacteria remains unknown, our results demonstrate that the two size classes of bacteria responded differentially to viral lysis of *Prochlorococcus*, resulting in succession between the large and small bacterial classes.

### The temporal nature of enhanced virus production in the presence of the grazer

The temporal nature of enhanced virus production led us to hypothesize that viruses were released earlier in the presence of the grazer. To assess this possibility, we measured the amount of virus DNA produced within *Prochlorococcus* cells (intracellular DNA) versus the amount released into the extracellular medium during cell lysis, whether encapsidated in virus particles or as unpackaged free virus DNA (extracellular DNA), over the first 30 hours of the experiment. At 12 h there was no difference in overall total virus DNA (intracellular+extracellular) in the presence or absence of the grazer (Fig. 3, Fig. S5) (RM ANOVA; p=0.613). However, the partitioning of DNA differed, with an average of 2.7-fold less virus DNA within *Prochlorococcus* cells (p=0.023) and 3.9-fold more virus DNA in the extracellular medium (p=0.013) in the combined grazer and virus treatment relative to the virus only treatment (Fig. 3, Fig. S5). In addition, large increases in cellular compounds such as glutamic acid and *Prochlorococcus* DNA were observed earlier in the extracellular medium in the low-grazer-high-virus treatment (12-18 h) relative to the high-virus treatment (18-24 h) (Fig. S4c,d). Together these findings provide support for earlier cell lysis and virus release in the presence of the grazer. This would subsequently allow a new cycle of infection to begin earlier and for overall greater production of viral DNA (Fig. S5 for all three replicates) and viruses (Fig. 1d) at 18 h in the presence of the grazer.

Earlier virus release in the presence of the grazer could result via several mechanisms. First, more rapid attachment could occur as a result of grazer-induced changes in the cell surface of *Prochlorococcus*. Second, grazers could induce changes in *Prochlorococcus* physiology, potentially through grazer release of material utilized by *Prochlorococcus*, which could result in an intrinsically shorter infection cycle. Third, since there is a lag between the intracellular formation of mature viruses and their release from the cell (4), egestion of fully formed viruses after grazer feeding on infected cells could lead to early release of viruses. Undigested and infectious viruses have been reported inside of, or egested from, protistan vacuoles (5, 6) as well as released from other zooplankton (7).

### Maintenance of grazer growth through utilization of other food sources in the system

Our findings of similar grazer growth in the presence of the virus despite reduced *Prochlorococcus* mortality (Fig. 2, Fig. S3) raised the question of whether grazer populations were being maintained by acquiring energy from sources other than healthy *Prochlorococcus* cells. Grazers may feed directly on viruses, as is known for some grazer-virus pairs (8–10). To assess if *P. bandaiensis* can feed on the P-SSP7 cyanophage, we ran additional treatments consisting primarily of the grazer and the virus (after depletion of *Prochlorococcus*). No decline in virus numbers was observed nor was there an increase in grazer abundance (Fig. S6). Thus, *P. bandaiensis* did not feed directly, nor grow, on the P-SSP7 cyanophage.

Other food sources for the grazers, as described in the main text, are the co-occurring heterotrophic bacteria whose growth was stimulated by organic matter released from *Prochlorococcus* as a result of viral lysis (Fig. 1e). While an increase in bacteria in the grazer treatments was not observed, likely due to immediate grazing, grazer egestion and excretion also are likely to have stimulated bacterial growth, providing another feedback loop for organic matter initially derived from *Prochlorococcus*.

Furthermore, grazers likely obtained resources by feeding on infected and damaged cells produced by viruses and on aggregates and cell debris generated in the combined grazer-virus treatment. Damaged *Prochlorococcus* cells could serve as a source of high-density nutrition for the grazers since these damaged cells consist mainly of membrane and cell wall components high in lipids and proteins. Furthermore, damaged cells without the low-density nutrition of the cytoplasm are smaller than healthy cells, potentially allowing for packing more resources of high nutritional value into the feeding vacuoles of the grazer.

### *Buoyancy of* Prochlorococcus *and infected* Prochlorococcus *cells*

Based on its small size, *Prochlorococcus* is generally assumed to be neutrally buoyant, and thus suspended in the water column (11, 12). Virus infection could affect the buoyancy of infected cells. We experimentally assessed the buoyancy of *Prochlorococcus* using settling columns. *Prochlorococcus* was indeed neutrally buoyant, with no significant differences in *Prochlorococcus* abundance between the settling column compartments (Fig. S11a). Infected *Prochlorococcus* cells were positively buoyant, and thus float, as seen from fewer infected cells in the bottom compartment relative to the top (p=0.0057) and middle (p=0.0096) compartments (Fig. S11b). Increased buoyancy is likely due to the reduction in sucrose and other osmolytes in infected *Prochlorococcus* cells (3).

## Supplementary Materials and Methods

### Quantification of intracellular and extracellular

Prochlorococcus *and virus DNA* Intracellular virus DNA was quantified using a heat lysis method (4, 13, 14). Briefly, cells from 1 mL were collected on 0.2 µm polycarbonate filters and washed three times with sterile seawater and once in preservation solution (10 mM Tris pH8, 100 mM EDTA, 0.5 M NaCl). Filters were flash frozen in liquid nitrogen and stored at -80 °C. Prior to DNA extraction, filters were immersed in 10 mM Tris (pH8) and cells removed by agitation in a Mini-Beadbeater (Biospec Products) for 2 min at 5000 rpm without beads. The cells were then heated at 95 °C for 5 min to lyse the cells and extract the DNA. Virus DNA was then quantified using SYBR green qPCR assays as described above.

Samples for extracellular *Prochlorococcus* and virus DNA were filtered through 0.22 µm syringe filters (see above). The filtrate (untreated) was diluted 100-fold in 10 mM Tris buffer (pH8) and subjected to SYBR green qPCR assays. Virus DNA was quantified using the same qPCR conditions and primers as described above. This consisted of both encapsidated and unencapsidated virus DNA. *Prochlorococcus* DNA was quantified using the same PCR thermocycling conditions, but with ITS primers (Forward: TACCTCCACTGAATACCACCTCT; Reverse: CGCACAAATAATAAATCTGCATCAT (15)) and serial dilutions of *Prochlorococcus* MED4 genomic DNA for standard curves.

### Organic matter quantification

Dissolved glutamic acid: Samples for quantification of dissolved glutamic acid in the extracellular medium were collected by filtering 40 mL of culture media through a 0.2 µm pore size Omnipore filter using gentle vacuum filtration. The filtrate was frozen and stored in acid washed 50 mL polypropylene falcon tubes at -20°C until analysis. Dissolved glutamic acid was extracted from the media using cation-exchange solid phase extraction (16) and measured using liquid-chromatography-mass spectrometry in which HILIC chromatography was coupled to a Thermo QExactive HF mass spectrometer (17). Chromatograms were integrated in Skyline (18) and best-matched internal standard normalization was employed to normalize for instrument variability (17). Glutamic acid was quantified by comparison to an authentic standard and adjusted for extraction efficiency, following Sacks et al. (16).

Particulate and dissolved ATP: Samples for particulate ATP (3x50 mL subsamples), including media blanks, were filtered onto GF/F filters. Particulate ATP was extracted from the filters by adding them to boiling 20 mM Tris buffer (pH7.4) for 5 min. The filtrate from the three subsamples was pooled (150 mL in total) for analysis of dissolved ATP. Both the particulate and dissolved ATP samples were frozen and stored at 20 °C until analysis. Dissolved ATP samples were collected from the control, low-virus and high-virus treatments only. Duplicated 50 mL subsamples were amended with 5 mM NaOH to initiate brucite formation that co-precipitates dissolved ATP (19, 20). Samples were centrifuged (1000Xg) for 1 h and the pellet was dissolved in 50 µL of 2.5 N HCl and brought up to 0.5 mL in deionized water. ATP concentrations were analyzed using the firefly bioluminescence method (21) on a Turner Biosystems luminometer (TBS2020^n^). Firefly lantern extract (FLE-250, Sigma-Aldrich) was diluted with equal parts of magnesium sulfate (0.04 M) and sodium arsenate buffers (0.1 M, pH 7.4). The concentration of ATP was determined using standard curves of known amounts of ATP.

Free DNA: Free DNA in the extracellular medium was determined from filtrates (5 mL) collected after filtration through GF/F filters. This fraction contains free DNA from all organisms in the extracellular medium as well as viral DNA inside capsids. Therefore, we determined both DNA in the entire fraction (total DNA), as well as the amount of DNA remaining after DNase treatment (DNase treatment is as described in the section on virus quantification (22)), which would degrade free unencapsidated DNA. Extracellular DNA was then determined following Brum et al. (23). Total DNA and the DNA remaining after DNase treatment was concentrated using centrifugal ultrafiltration with 10 kDa cut-off centrifugal filters (Amicon). The medium was exchanged for a Tris-EDTA buffer (10 mM Tris-HCl, 1 mM EDTA, pH8) and concentrated to ∼0.1 mL. DNA was then quantified by fluorescence using the Qubit HS dsDNA Assay kit (ThermoFisher Scientific). Free DNA was calculated by subtracting the amount of DNA remaining after DNase treatment from the total DNA in the GF/F filtrate.

### Settling experiments

Settling experiments were carried out for triplicate control, and duplicate high-virus, high-grazer and high-virus-high-grazer treatments using subsamples collected from the large carboys at T=12 and T=24 h and incubated for 4 h in the settling columns under the growth conditions described above using the SETCOL method (24). We measured the general *Prochlorococcus* population and infected *Prochlorococcus* cells in the three compartments of the settling column to distinguish positively, neutrally or negatively buoyant cells at the end of the 4 h incubation.

*Prochlorococcus* abundances were quantified using flow cytometry as described above. Infected *Prochlorococcus* cells were quantified using the iPolony method, a solid phase PCR method (25). Sorted *Prochlorococcus* cells were subjected to solid phase PCR using specific primers and probe for the P-SSP7 cyanophage (Forward: TAACGCAACACAAGGATACAGAGAA, Reverse: ACCTCCGTCATATCTGCCTAAGTAA, Probe: GGTATGACTATGGTTGGAGCGGATT (22) to determine the percentage of cells that were infected by the P-SSP7 phage in each compartment. Free viruses were quantified in the filtrate of the sorted samples and subtracted to determine cell infection.

### Prochlorococcus mortality estimates

The *Prochlorococcus* mortality rate induced by a specific treatment (grazing, viral lysis or both) was quantified by calculating the difference between the net growth rate in the control treatment and the treatment of interest. Our approach assumed that per-capita division rates were not altered by the predator/virus manipulations, such that treatment differences in net growth reflect differences in loss rates (cf. dilution-based formulations of *μnet=μ−loss*) (26). Net growth rates were calculated from the abundance data as the slope of the natural logarithm of (smoothed) abundance over time (i.e., i(*t*)≈Δln *N*/Δ*t*), to linearize exponential growth, improving growth dynamics analysis. Smoothing splines were used to reduce high-frequency sampling noise while preserving the underlying dynamics. Treatment-induced mortality (loss) was then calculated as the difference between net growth rates in the control and the treatment, interpreting the control as the baseline with virtually no loss.

We then developed models of grazer-induced and virus-induced mortality based on the relationship between prey/host abundance and consumption rates. To relate prey/host abundance to per-capita grazing and viral lysis rates, we compared two standard functional-response forms, sensu Holling (27): a Holling type I response *f(N)=aN*, where *f(N)* is the per-capita ingestion or lysis rate, *N* is prey/host abundance, and *a* is the clearance/attack coefficient describing how consumption increases with prey/host density; and a Holling type II response *f(N)=aN / (1+ahN)*, where *h* is the handling time per prey item (which implies an asymptotic maximum rate *fmax =1/h* at high *N*). Both models were fitted by minimizing the root mean squared error (RMSE) between observed and predicted per-capita rates using Differential Evolution optimization (DEoptim, (28)). Search bounds were set to [0, 1000] for all parameters, with convergence tolerance 10⁻⁶ and a maximum of 1000 generations. Model performance was assessed using RMSE, the coefficient of determination (R²), and the Akaike Information Criterion (AIC), which penalizes the additional parameter in the type II model.

For grazing on *Prochlorococcus*, the type I model yielded a = 56.8 cells grazer⁻¹ h⁻¹ (10⁷ cells mL⁻¹)⁻¹ (R² = 0.921, RMSE = 6.90, AIC = 342). The type II model improved the AIC by 14 units (AIC = 328, R² = 0.934, RMSE = 6.30), but converged to h = 0.005, implying a half-saturation constant of ∼2.6 × 10⁷ cells mL⁻¹, roughly twice the highest prey abundance observed. Because the AIC difference favors type II, this model was used for grazer predictions, while noting that the two models produced nearly identical predictions across the observed range of host (Fig. S12a). For viral lysis, the type I model yielded a = 5.18 × 10⁻³ cells phage⁻¹ h⁻¹ (10⁷ cells mL⁻¹)⁻¹ (R² = 0.814, RMSE = 1.71 × 10⁻³, AIC = −699). The type II optimizer converged to h ≈ 0, recovering the type I solution (RMSE = 1.71 × 10⁻³, AIC = −697, ΔAIC = +2). The type II model thus provided no biologically meaningful evidence of prey saturation within the measured range (Fig. S12b). Therefore, we retained the simpler type I model for virus-induced mortality predictions. Parameter uncertainty was estimated by nonparametric bootstrap resampling (1000 iterations), refitting the type I or type II model to each resample. The 95% confidence intervals are shown in (Fig. S12).

The fitted models were then used to predict mortality rates under experimental conditions. These predicted rates were multiplied by the corresponding grazer or virus abundance to estimate predicted loss rates due to grazing or viral lysis; for the combined treatment, we calculated the total mortality rate as the sum of the predicted loss rates induced by both virus and grazer, assuming independent and additive effects (29).

## List of Supplementary Tables

**Table S1:** Starting abundances of *Prochlorococcus*, cyanophage, grazer, and co-occurring heterotrophic bacteria.

| Treatment | *Prochlorococcus*<br>MED4<sup>a</sup><br>(cells mL<sup>-1</sup>) | Cyanophage<br>P-SSP7<sup>b</sup><br>(pfu mL<sup>-1</sup>) | *Paraphysomonas*<br>*bandaiensis*<sup>a</sup><br>(cells mL<sup>-1</sup>) | Heterotrophic<br>bacteria<sup>a</sup><br>(cells mL<sup>-1</sup>) |
| --- | --- | --- | --- | --- |
| **Control** | 1.71±0.08×10<sup>7</sup> | - | - | 0.96±0.21×10<sup>6</sup> |
| **Low Virus** | 1.77±0.05×10<sup>7</sup> | 2.40×10<sup>4</sup> | - | 0.85±0.31×10<sup>6</sup> |
| **High Virus** | 1.64±0.04×10<sup>7</sup> | 1.20×10<sup>7</sup> | - | 0.96±0.36×10<sup>6</sup> |
| **Low Grazer High Virus** | 1.76±0.12×10<sup>7</sup> | 1.20×10<sup>7</sup> | 6.01±2.07×10<sup>2</sup> | 0.68±0.14×10<sup>6</sup> |
| **Low Grazer** | 1.47±0.14×10<sup>7</sup> | - | 6.23±0.85×10<sup>2</sup> | 1.18±0.07×10<sup>6</sup> |
| **High Grazer** | 1.63±0.10×10<sup>7</sup> | - | 4.68±0.25×10<sup>3</sup> | 1.47±0.49×10<sup>6</sup> |

<sup>a</sup>Average and standard deviation of 3 biological replicates except for the control which is the average of 4 biological replicates.
<sup>b</sup>The titer of the phage added to the treatments 1.5 h prior to the beginning of the experiments.

**Table S2.** Statistical analyses. (excel table)

**Table S3.** Change point analysis of damaged *Prochlorococcus* cells and fluorescent aggregates for the replicates of the different treatments. (excel table)

**Table S4.** Source of SEM images appearing in the manuscript. (excel table)

**Figure S1.**
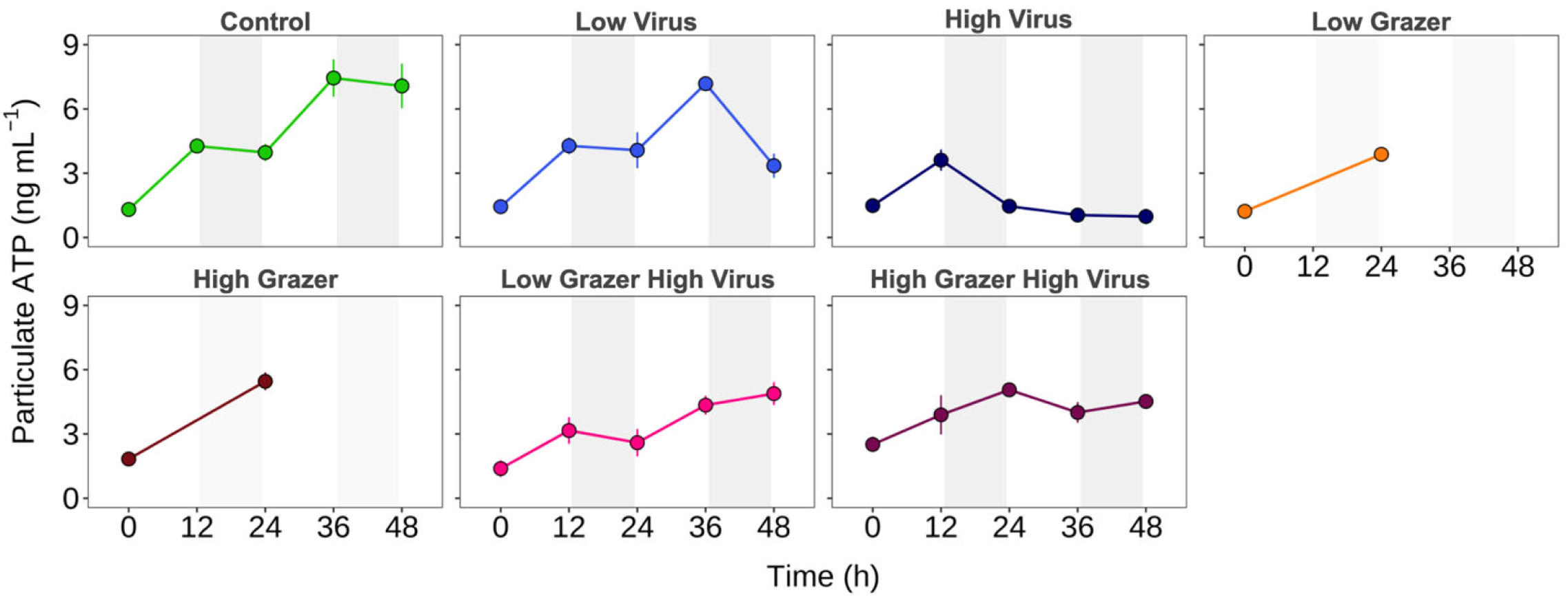
Concentrations of particulate ATP as a proxy for cell biomass. Particulate ATP in the different treatments derived from all organisms collected on GF/F filters, which include *Prochlorococcus,* grazers (where relevant), heterotrophic bacteria and aggregates. Shading indicates nighttime. Average and standard deviation of 3 biological replicates. Fewer samples were available for analysis for the low-grazer and high-grazer treatments due to technical constraints. See Table S2L for statistical analyses.

**Figure S2.**
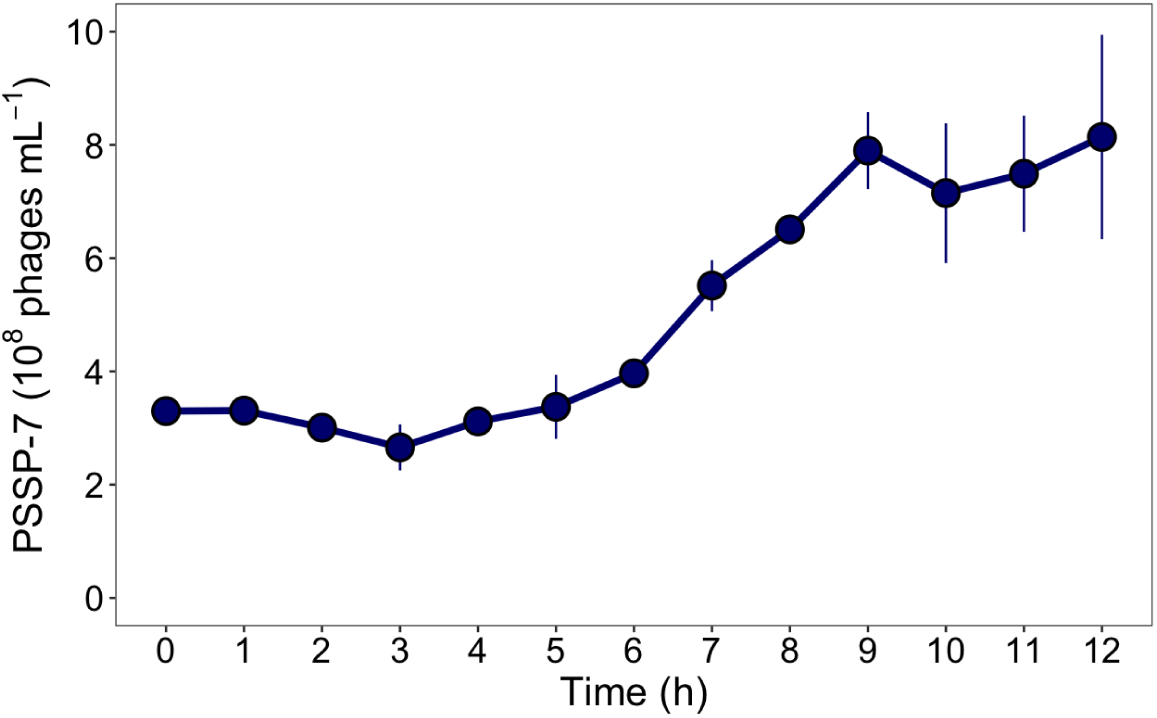
Single-step growth curve of the P-SSP7 cyanophage under conditions used in this study. Exponentially growing *Prochlorococcus* MED4 at 50 µmol photon m^-1^ s^-1^ for 14 h light and 10 h dark at 21 °C with gentle stirring was infected with P-SSP7 at a multiplicity of infection of 3. Infective phage release was measured by the plaque assay at different times after cyanophage addition, showing that the length of the lytic cycle was between 6-9 hours. Average and standard deviation of 3 biological replicates. Infection was done entirely during the light period.

**Figure S3.**
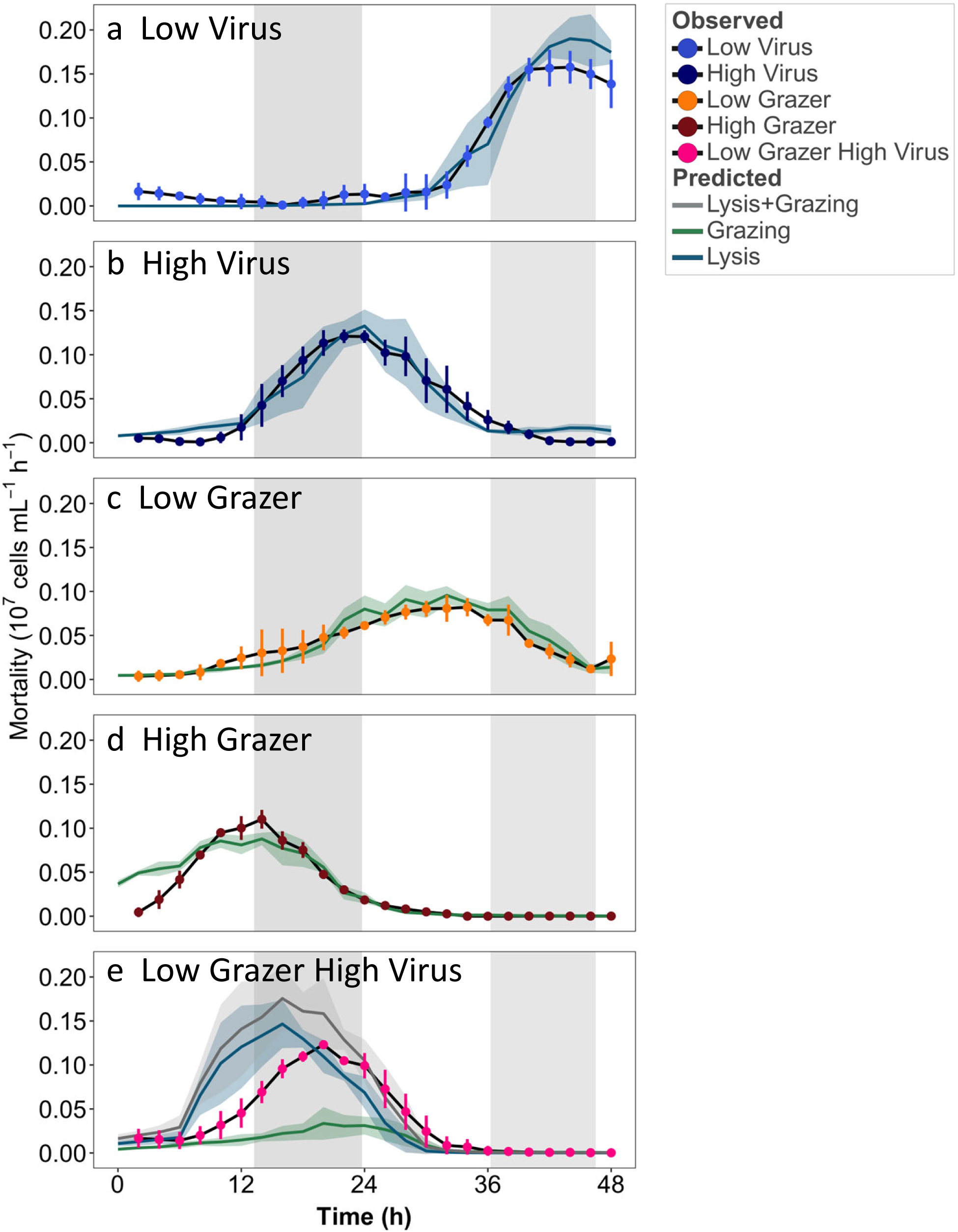
*Prochlorococcus* mortality estimates. Observed mortality rates (black lines and colored circles) and predicted mortality rates (lines without symbols) of *Prochlorococcus* in the low-virus (a), high-virus (b), low-grazer (c), high-grazer (d), and low-grazer-high-virus (e) treatments. Observed mortality was calculated based on the difference in net growth rates between the control (assumed to have virtually no loss) and the treatment of interest. Predicted mortality rates are based on a model that predicts grazing rates and viral lysis based on the functional responses of grazers and viruses to *Prochlorococcus* abundance. Vertical bars and the shaded areas surrounding the lines represent the standard deviation of 3 biological replicates. The blue and green lines in (e) indicate predicted mortality rates due to viral lysis and grazing, respectively, while the grey line indicates the total predicted mortality, calculated as the sum of mortality due to lysis and grazing. Vertical shading indicates nighttime. Panel (e) is the same as Fig. 2 and is presented here for ease of comparison.

**Figure S4.**
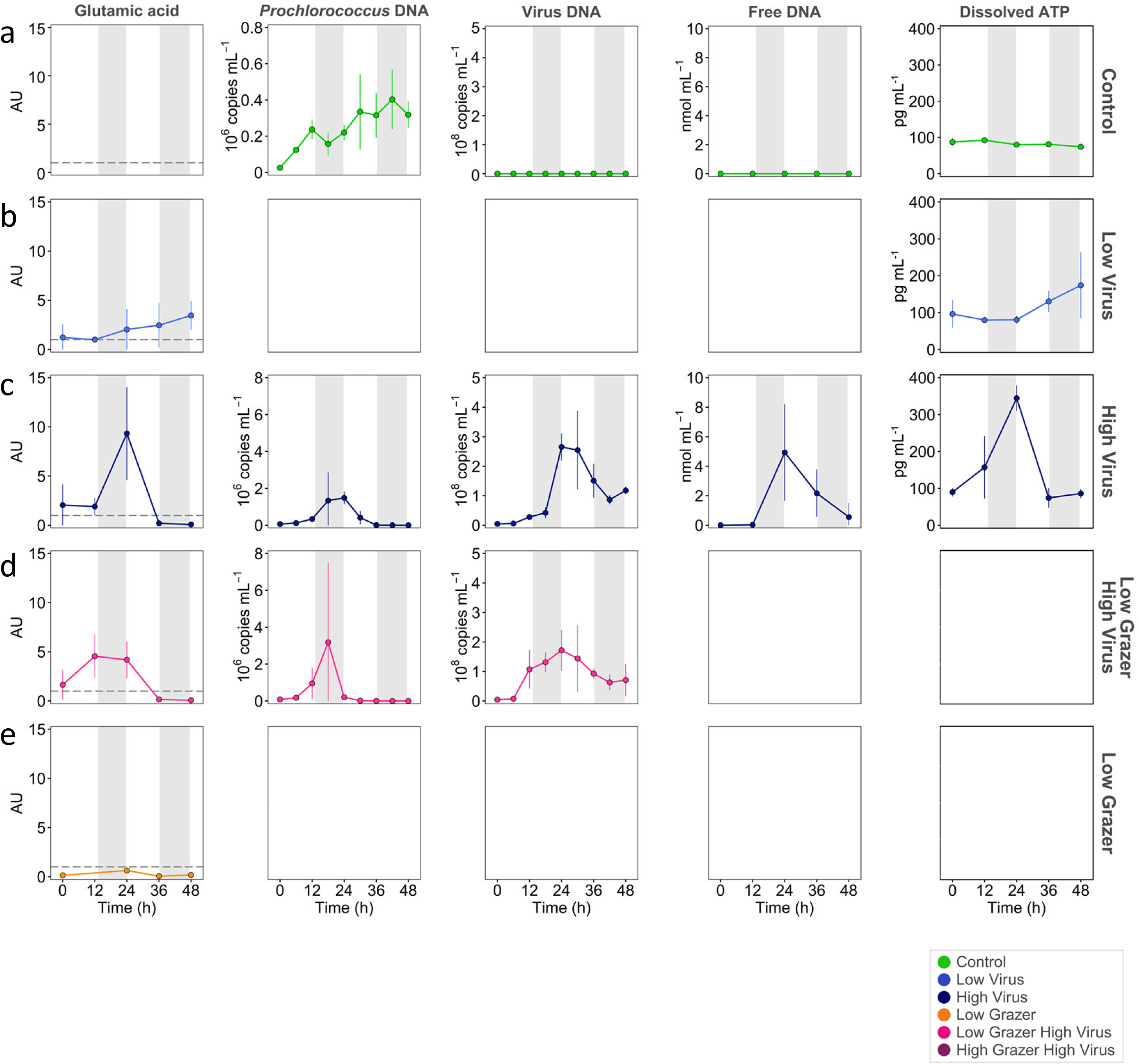
Organic matter in the extracellular medium. Glutamic acid, *Prochlorococcus* DNA, virus DNA, free DNA, ATP in the extracellular medium of the control (a), low-virus (b), high-virus (c), low-grazer-high-virus (d), and low-grazer (e) treatments. Note the 10-fold lower scale for *Prochlorococcus* DNA in the control (a) relative to the high-virus and low-grazer-high-virus treatments (b,c). Glutamic acid levels were normalized to the average of that in the control treatment. Glutamic acid levels were normalized to the average of that in the control treatment, which was set to 1 and is shown as a dashed line, including for the control (a). Note that not all compounds were analyzed for all treatments, hence the empty boxes. *Prochlorococcus* and virus DNA were determined by qPCR, whereas free DNA from all biological sources was determined using a DNA fluorescence assay. Shading indicates nighttime. See Table S2M,N,O,P,Q for statistical analyses.

**Figure S5.**
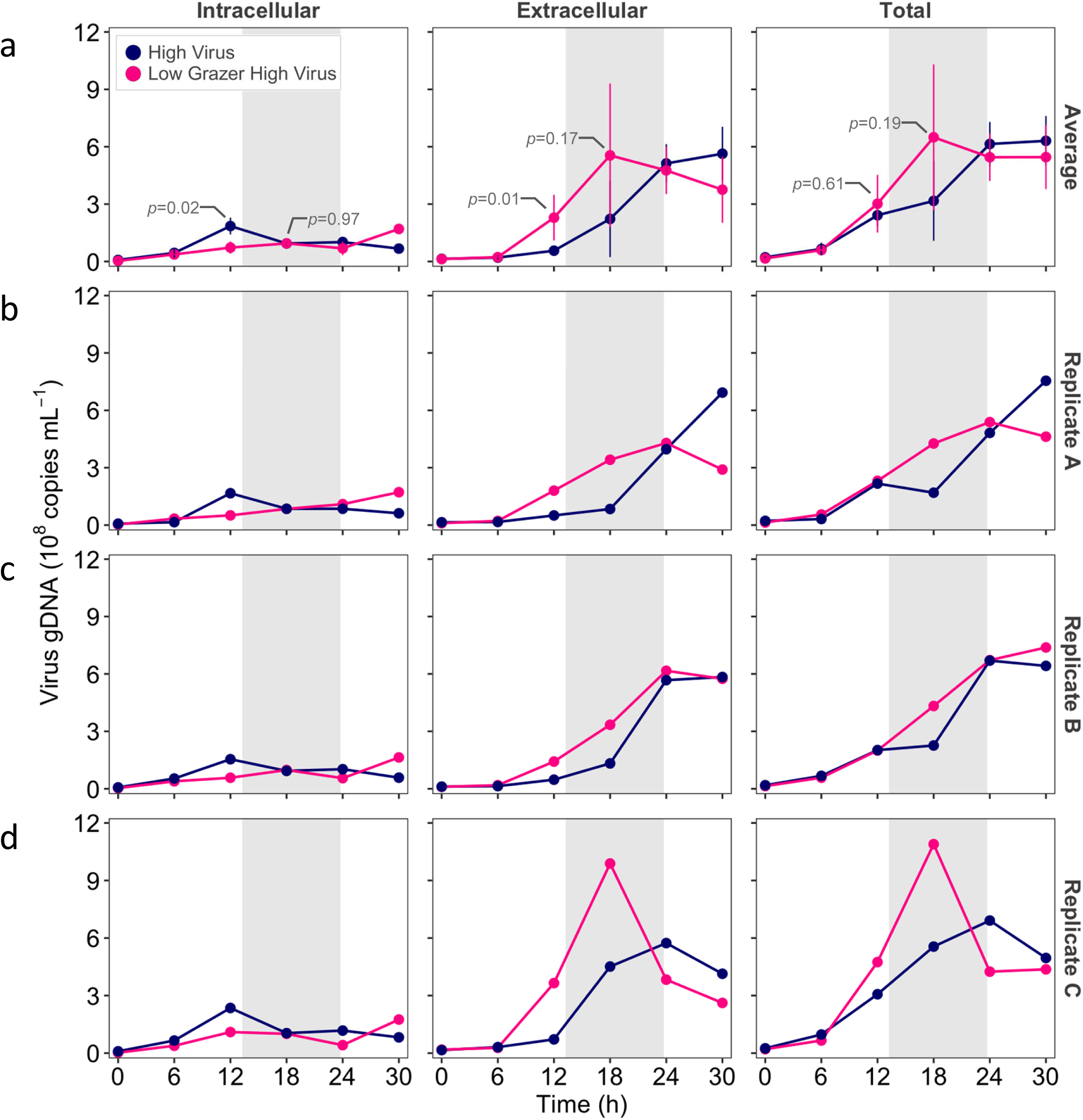
Partitioning of virus DNA in intracellular or extracellular fractions for the average and individual replicates. Intracellular virus DNA (left panel), extracellular virus DNA (middle panel), and total intracellular and extracellular DNA (right panel) during infection of *Prochlorococcus* MED4 in the absence (blue) and presence (pink) of the grazer. (a) Average and standard deviation of 3 biological replicates, (b) Replicate A, (c) replicate B, and (d) replicate C. Extracellular DNA includes encapsidated DNA packaged inside virus particles as well as unpackaged free DNA released from cells during lysis. Repeated measure analysis on log transformed data indicates a significant difference between the treatments. Post-hoc Tukey p-values for 12 and 18 h are shown. While p-values were greater than 0.05 at 18 h for extracellular and total DNA (a), this is likely due to the variability in values between replicates since greater values were found in the combined grazer-virus treatment for all three replicates (b,c,d). Shading indicates nighttime. See Table S2I,J,K for statistical analyses. The average data shown in (a) are the same as Fig. 3 and are shown here for ease of comparison to the individual replicates (b,c,d).

**Figure S6.**
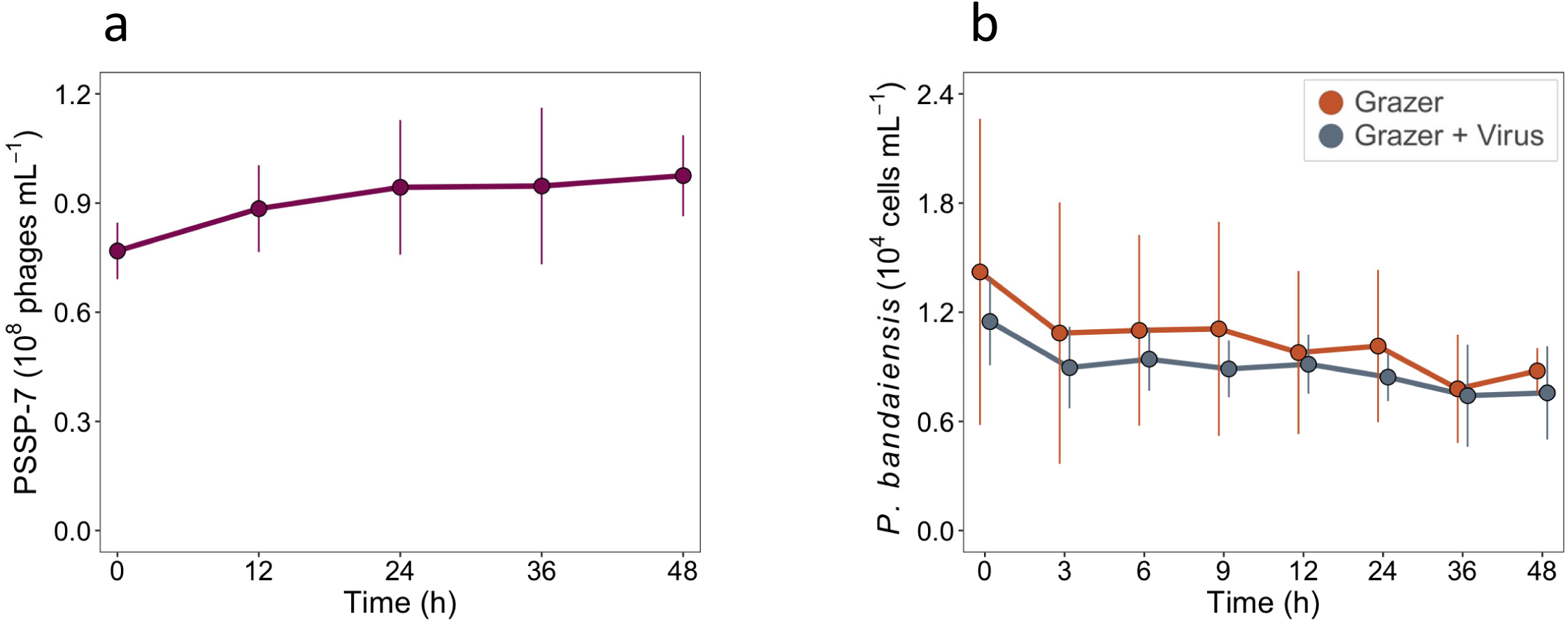
Virus and grazer abundances in the absence of *Prochlorococcus* and heterotrophic bacteria. Abundances of the P-SSP7 phage in the presence of the grazer alone (a), and abundances of the *P. bandaiensis* grazer (b) when alone (orange) or with viruses (grey), but without virtually any *Prochlorococcus* or bacterial cells (below 1.8×10^4^ *Prochlorococcus*·mL^-1^ and 4.1×10^4^ heterotrophic bacteria·mL^-1^ at T=0 h). Average and standard deviation of 3 biological replicates.

**Figure S7.**
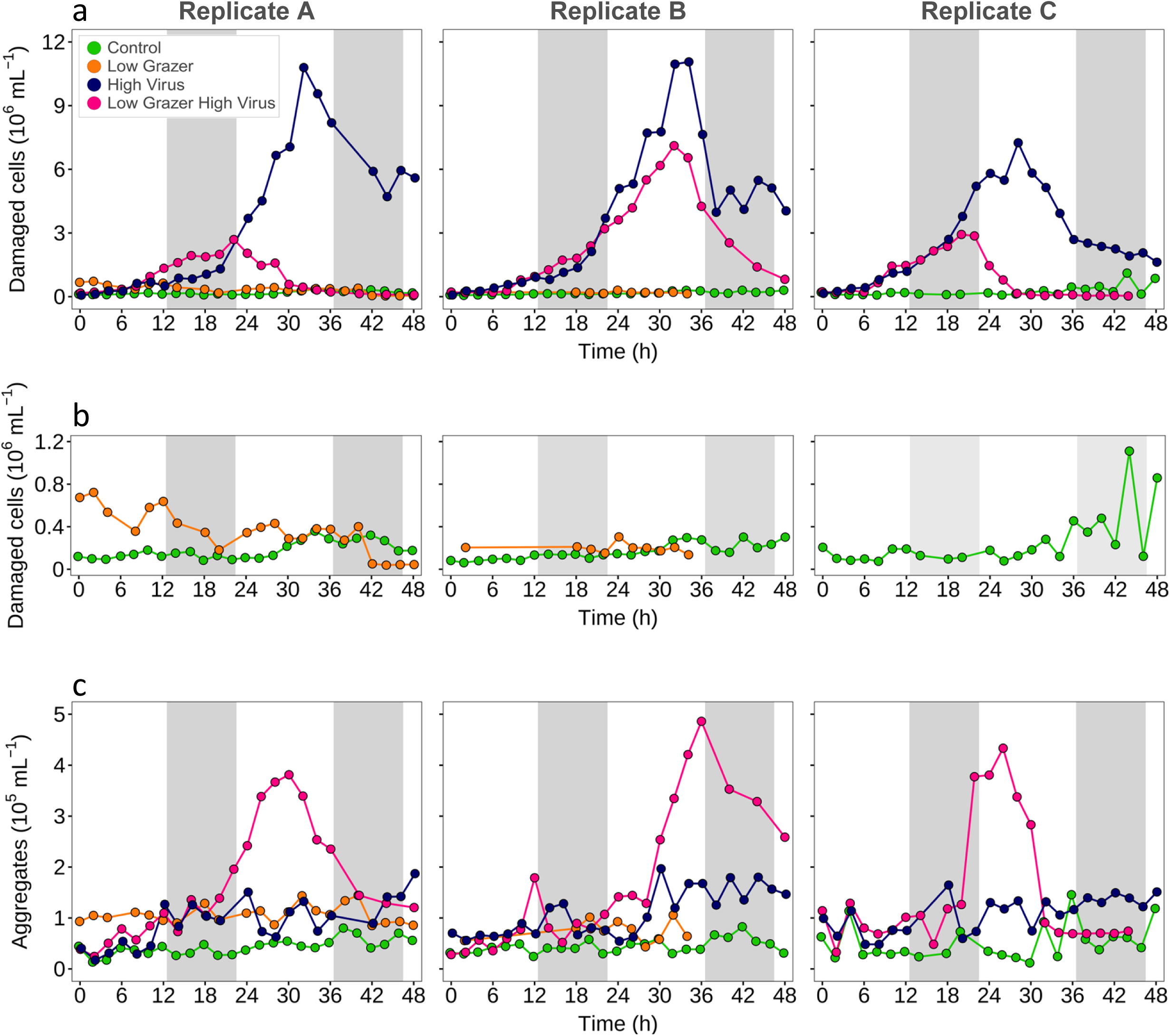
Appearance of damaged *Prochlorococcus* cells and fluorescent aggregates in individual replicates. Abundances of damaged *Prochlorococcus* cells (low-scatter population) (a), zoomed scale of damaged cells (b), and fluorescent aggregates (intermediate-scatter population) (c), in replicate A (left panels), replicate B (middle panels) and replicate C (right panels), for the control, low-grazer, high-virus and low-grazer-high-virus treatments (a, c), and in the zoomed-in view in the control and low grazer treatments (b). Due to technical issues, data from the low grazer treatment is missing for replicate C and part of replicate B. The results for replicate A are the same as those shown in Fig. 4. Shading indicates nighttime. See Table S3 for the timing of treatment-specific changes in damaged cells and fluorescent aggregates and Table S2G,H for statistical analyses.

**Figure S8.**
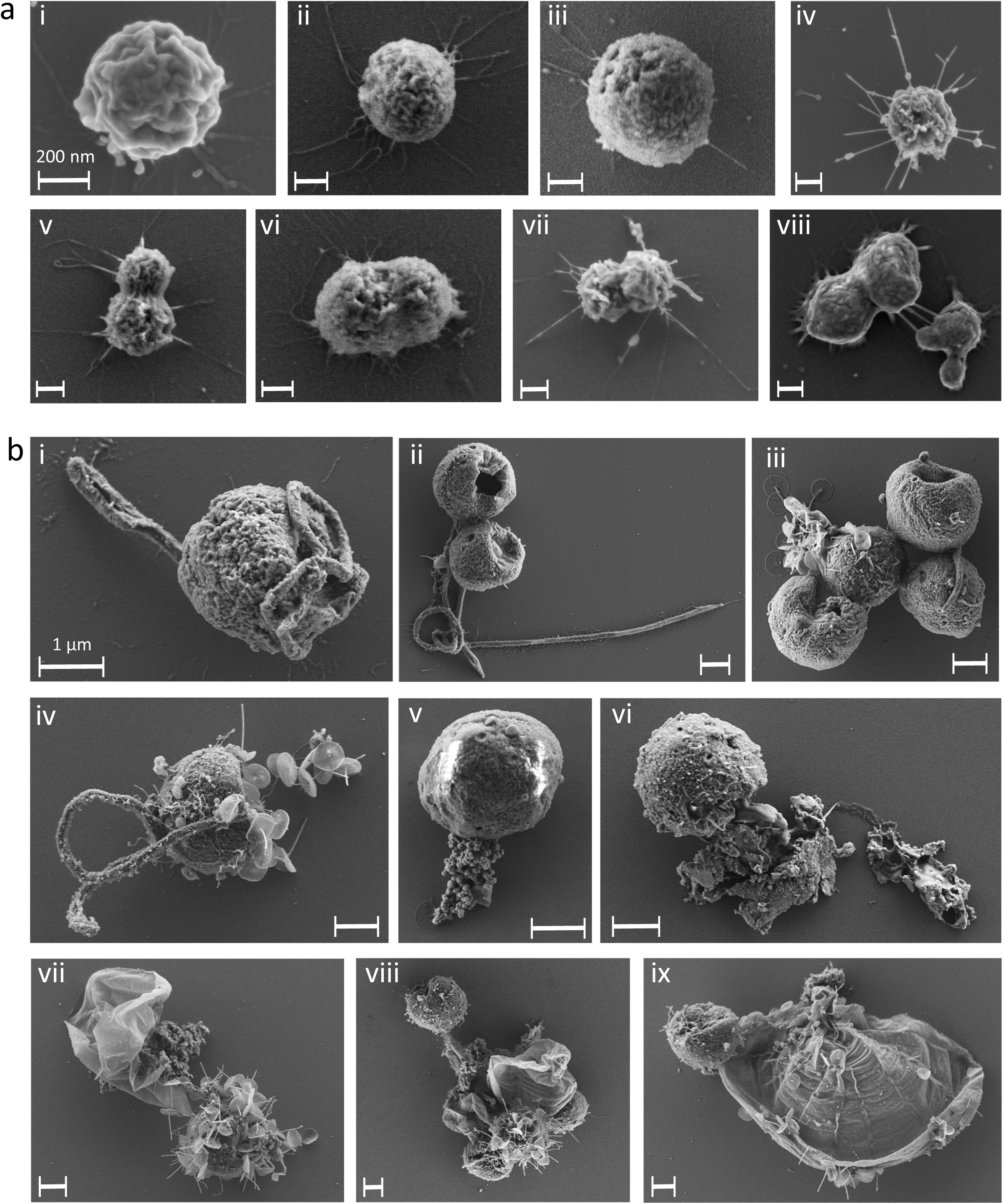
SEM images of *Prochlorococcus* and grazers. (a) Representative SEM images of intact *Prochlorococcus* cells from the control treatment at T=32 h (a), and grazers from the low-grazer-high-virus treatment at T=32 or 34 h (b). The scale bar, shown in white, indicates a length of 200 nm for images in (a) and 1 µm for images in (b). See Table S4 for the source of each image with respect to biological replicate and time point.

**Figure S9.**
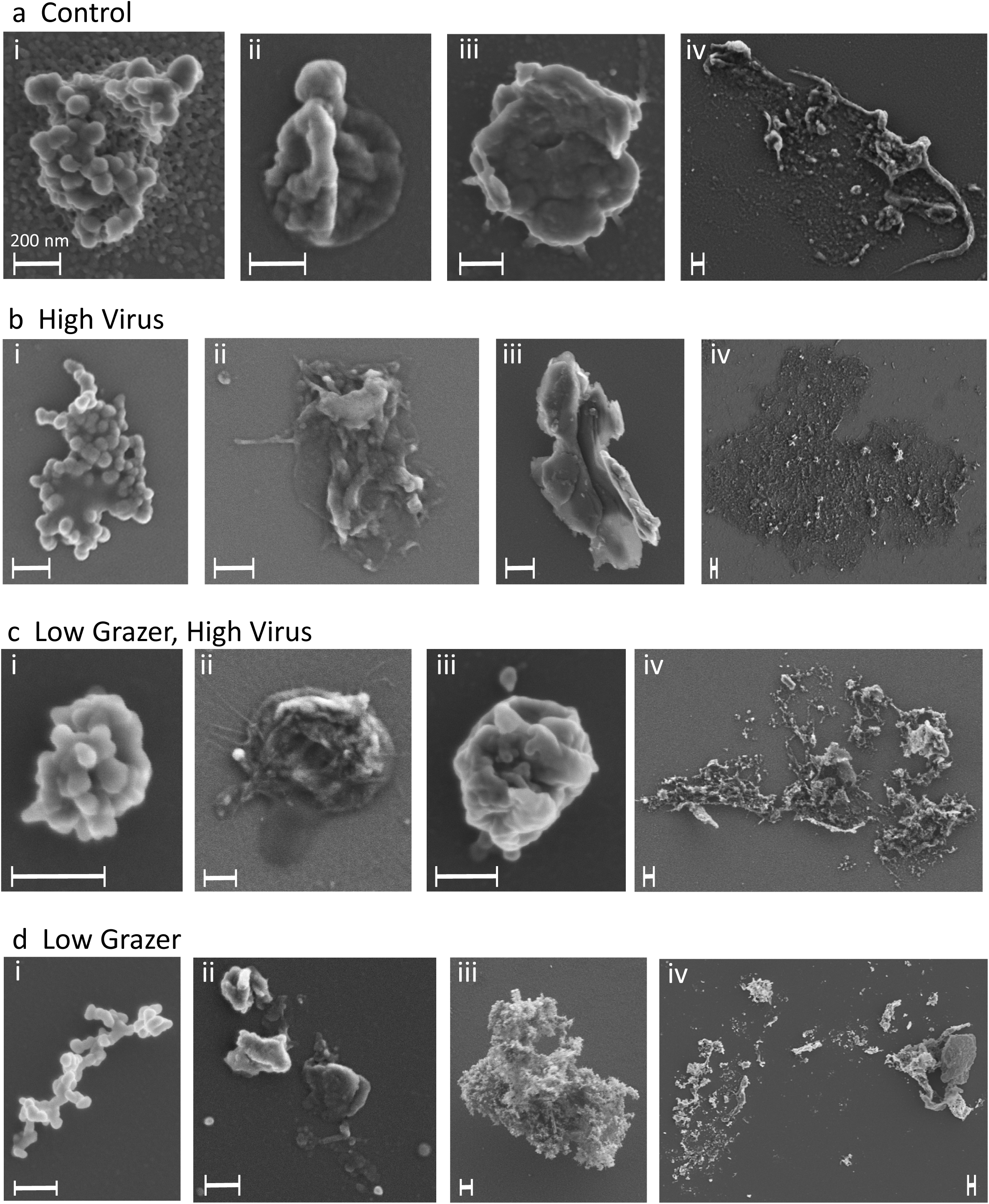
SEM images of damaged cells. Representative SEM images of damaged cells and more diffuse cellular debris from the control with *Prochlorococcus* alone (a), and *Prochlorococcus* from the high-virus (b), low-grazer-high-virus (c), and low-grazer (d) treatments from T=32 or 38 h. The white scale bars are 200 nm in length in all images. See Table S4 for the source of each image with respect to biological replicate and time point.

**Figure S10.**
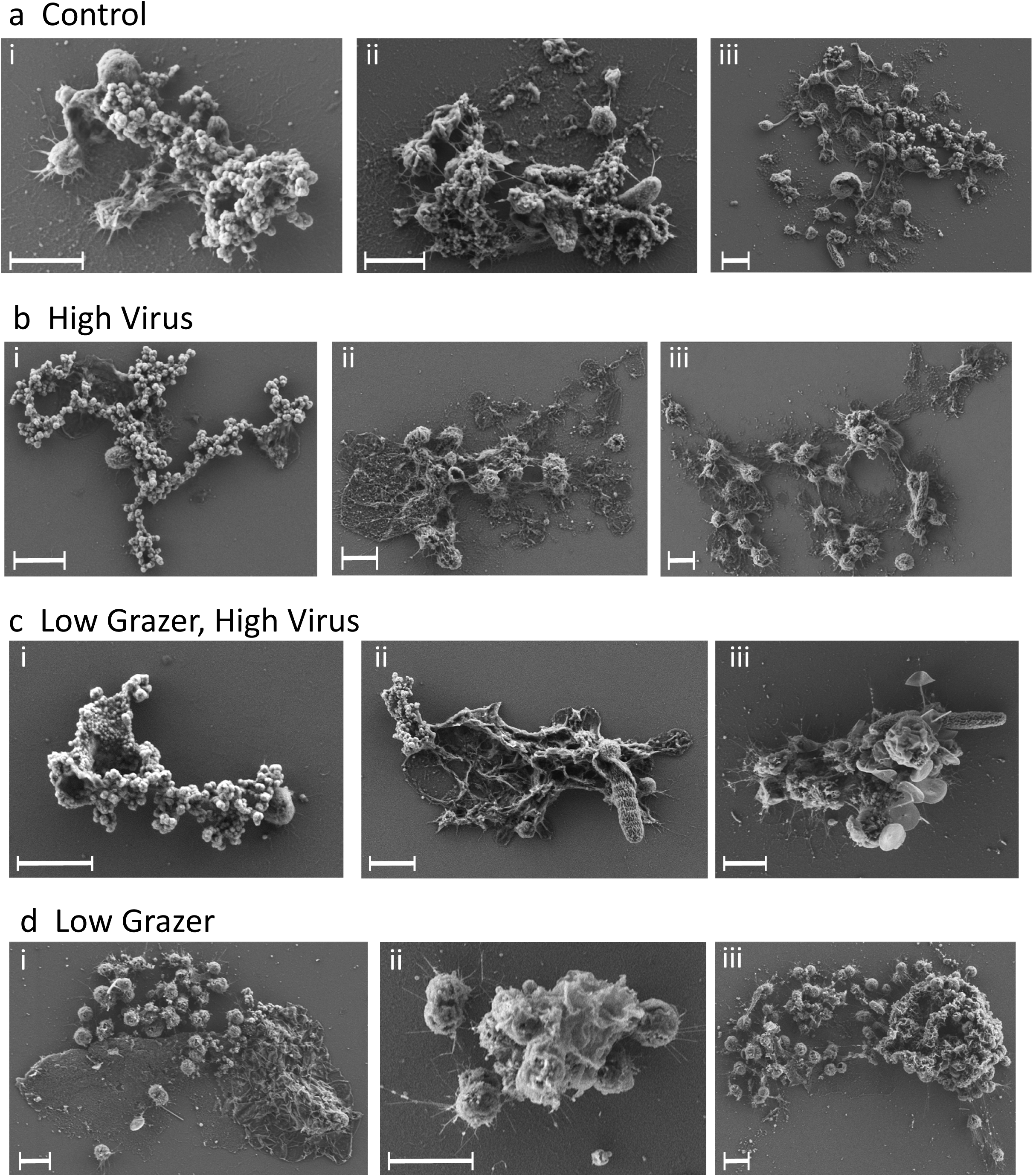
SEM images of fluorescent aggregates. Representative SEM images of fluorescent aggregates from the control with *Prochlorococcus* alone (a), and *Prochlorococcus* subjected to high-virus (b), low-grazer-high-virus (c), and low-grazer (d) treatments from T=32 or 38 h. The white scale bars are 1 µm in length in all images. See Table S4 for the source of each image with respect to biological replicate and time point.

**Figure S11.**
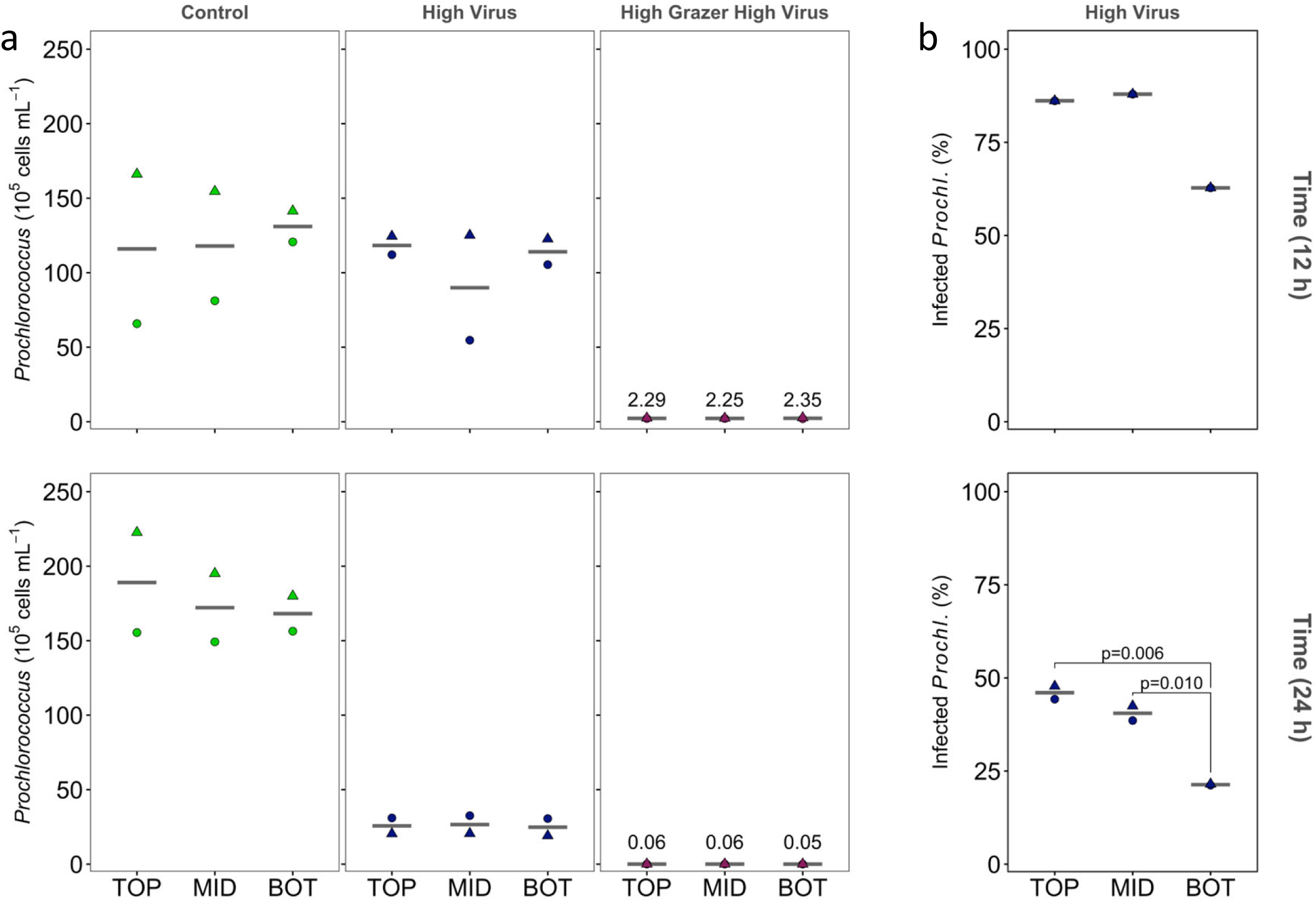
Buoyancy of healthy and infected *Prochlorococcus* cells. Partitioning within the top, middle or bottom compartments of the settling column of *Prochlorococcus* MED4 (a) and *Prochlorococcus* infected with P-SSP7 cyanophage (b), in the control, high-virus and high-grazer-high-virus treatments (a) or the high-virus treatment (b) at 12 and 24 h after the beginning of the experiment. TOP= top, MID=middle, BOT=bottom compartments of the settling column. Symbols indicate different biological replicates. No significant differences were found for *Prochlorococcus* in the different compartments at T=12 h and at T=24 h (a), indicative of neutral buoyancy. Significantly fewer infected *Prochlorococcus* cells were found at T=24 h in the high-virus treatment in the bottom compartment relative to the top (p=0.0057) and the middle (p=0.0096) compartments (n=3) (b), indicative of positive buoyancy. Only two replicates are available for infected cells at T=12 h, therefore no statistical analysis was possible. See Table S2R for all statistical analyses.

**Figure S12.**
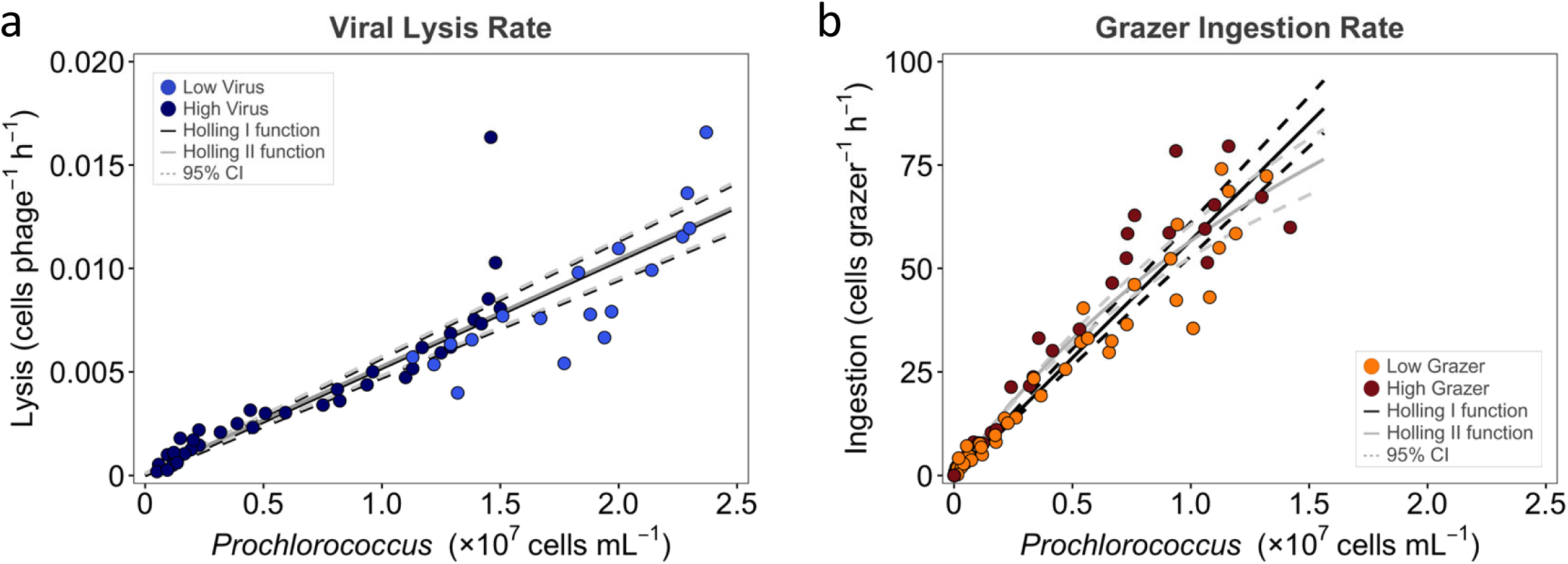
Functional-response fits for grazing and viral lysis of *Prochlorococcus*. (a) Estimated virus-induced lysis rate versus *Prochlorococcus* abundance in the low- and high-virus treatments. (b) Estimated grazer ingestion rate versus *Prochlorococcus* abundance in the low- and high-grazer treatments. Circles show observations in low- and high-grazer or virus treatments. Solid black lines show fitted Holling type I models (linear) and solid grey lines show fitted Holling type II models (hyperbolic). The dashed black and grey lines show the 95% confidence intervals (CI) for the Holling type I and type II models, respectively. The two model curves were virtually the same across the observed prey range (see Supplementary methods).

